# Identification of novel RNA Polymerase II CTD interaction sites on the mRNA Capping Enzyme

**DOI:** 10.1101/2020.02.25.964700

**Authors:** Marcus G. Bage, Rajaei Almohammed, Victoria H. Cowling, Andrei V. Pisliakov

## Abstract

Recruitment of the mRNA Capping Enzyme (CE/RNGTT) to the site of transcription is essential for the formation of the 5’ mRNA cap, which in turn ensures efficient transcription, splicing, polyadenylation, nuclear export and translation of mRNA in eukaryotic cells. The CE is recruited and activated by the Serine-5 phosphorylated carboxyl-terminal domain (CTD) of RNA polymerase II. Through the use of molecular dynamics simulations and enhanced sampling techniques, we provide a systematic and detailed characterisation of the human CE-CTD interface, describing the effect of the CTD phosphorylation state, length and orientation on this interaction. Our computational analyses identify novel CTD interaction sites on the human CE surface and quantify their relative contributions to CTD binding. We also identify differences in the CTD binding conformation when phosphorylated at either the Serine-2 or Serine-5 positions, thus providing insights into how the CE reads the CTD code. The computational findings are then validated by binding and activity assays. These novel CTD interaction sites are compared with cocrystal structures of the CE-CTD complex in different eukaryotic taxa, leading to the conclusion that this interface is considerably more conserved than previous structures have indicated.

## 2 Introduction

mRNA capping is an essential process required for efficient gene expression and regulation in all eukaryotic organisms (1). The mRNA cap prevents degradation by 5’-exonucleases during transcription and acts as a platform to recruit initiation factors required for splicing, polyadenylation, nuclear export and translation (2–8). mRNA is capped at the 5’-end with an inverted 7-methylguanosine moiety. This process occurs in three stages: i) the 5’-end triphosphate is hydrolysed to diphosphate; ii) GMP is covalently linked to the diphosphate 5’ end; iii) the guanosine base is methylated at the N7 position (1). In animals the first two stages are performed by a bifunctional protein, the Capping Enzyme (CE/RNGTT), which contains triphosphatase (TPase) and guanylyl-transferase (GTase) enzymatic domains separated by a disordered linker (9, 10). The mammalian CE GTase functions independently of the TPase domain (10–12). The final step, N7 methylation of the guanosine base, is performed by RNMT in complex with its activating mini-protein RAM (13, 14).

The process of mRNA capping is tightly coupled to transcription, occurring during the elongation phase (15, 16). At this stage the CE is recruited to the site of transcription by the RNA polymerase II (Pol II) carboxyl-terminal domain (CTD) (17, 18). The CTD is located in RPB1, the largest subunit of RNA Pol II, and is composed of a tandem repeated heptad motif with the consensus sequence Y_1_S_2_P_3_T_4_S_5_P_6_S_7_ (19, 20). This domain is disordered and can be dynamically phosphorylated at several positions to form a highly complex pattern known as the CTD phosphorylation code, which is used to recruit and regulate the transcription machinery, including the capping enzymes, at the correct phase of transcription (17, 18, 20, 21). Although each of the residues Tyr1, Ser2, Thr4, Ser5 and Ser7 can be phosphorylated and have all been shown to vary in their levels of phosphorylation during the transcription cycle, one fundamental transition occurs from the Ser5 to Ser2 phosphorylation state (pSer5 and pSer2) during transcription elongation (20, 22–24). The CE GTase domain is known to bind to the CTD during the elongation phase when the CTD is phosphorylated at the Ser5 position (12, 15, 16, 25). This localises the CE to the site of transcription and increases the rate of the first step of GTase catalysis. However, the importance of this activation effect on the regulation of mRNA capping remains unclear, with recent experiments indicating that the primary role of the GTase-CTD interaction is recruitment rather than allosteric activation (26, 27). Interestingly, the GTase can also bind to Ser2 phosphorylated CTD, however, this interaction does not stimulate GTase activity (12).

The CE GTase is highly conserved among eukaryotic organisms and is composed of three subdomains: i) the nucleotidyltransferase (NT) domain, which contains essential residues involved in catalysis, ii) the oligonucleotide-binding (OB) domain, predicted to bind mRNA for cap addition, and iii) the hinge domain that enables large-scale conformational changes to occur, opening and closing the active site to facilitate substrate binding, catalysis and product release (Figure 1A) (10, 28, 29). Three cocrystal structures of the CE GTase interacting with the Pol II CTD fragments were previously reported: one mouse GTase and two from yeast (*Candida albicans* and *Schizosaccharomyces pombe*) (30–32). Although the CTD binds to the NT domain in all of these structures, they display distinct CTD docking sites (CDSs). This has led to the conclusion that CTD recognition by the GTase is performed by distinct molecular mechanisms, with different taxa independently evolving different CTD interaction sites on the GTase surface (30, 32, 33).

**Figure 1.**
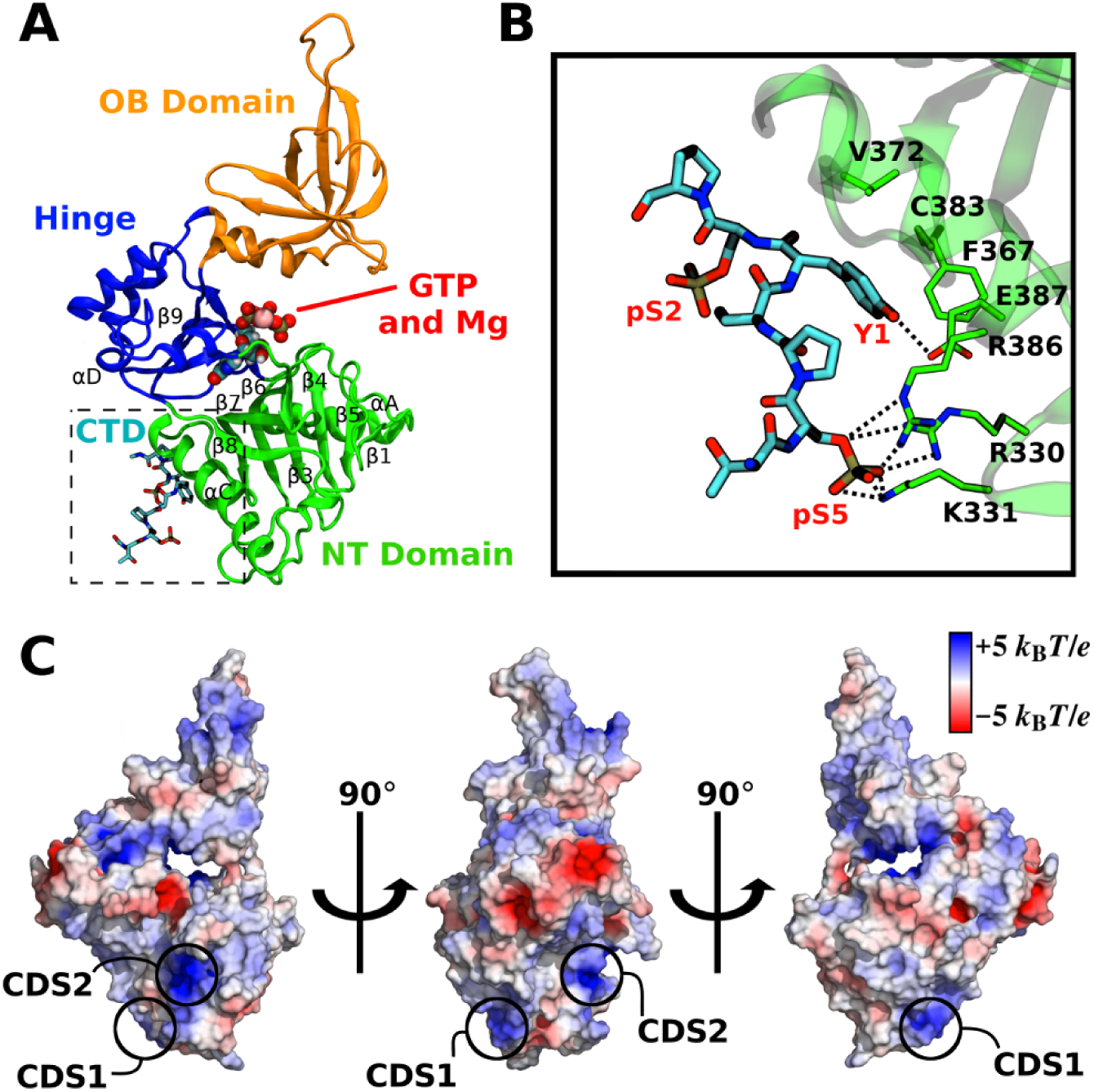
Structure of the Capping Enzyme GTase domain and its interactions with RNA Pol II CTD. (**A**) Structure of the human GTase domain (PDB ID: 3S24) (10) with one RNA Pol II CTD heptad (shown as sticks) bound in the conformation as resolved in the mouse GTase-CTD complex by Ghosh *et al.* (PDB ID: 3RTX) (32). Three subdomains of the GTase are labelled and coloured in green (NT), orange (OB) and blue (Hinge). GTP and Mg^2+^ (shown as spheres) were modelled in representative binding poses of the first enzymatic step and indicate the location of the active site. Important secondary structural elements are labelled following assignment in Chu *et al.* (10). (**B**) The GTase-CTD interface displaying the previously identified CTD interaction sites on the GTase: a pSer5 charged pocket (CDS1, composed of R330, K331 and R386) and a Tyr1 interaction site (CDS-Y1, composed of F367, V372, C383 and E387). The pSer2 group is solvent exposed and forms no interactions with the GTase residues. (**C**) Electrostatic potential surface of the human CE GTase. Positively charged regions (blue) have the potential to form additional pSer interaction sites. The pSer interaction sites discussed in this work—CDS1 and a novel site CDS2—are labelled.

The relatively low binding affinity of the CTD to the CE GTase (*K_d_* = 139 *µ*M) (32), in combination with the disordered and flexible nature of the CTD (34), the proposed CTD heptad looping out mechanism (31), and the GTase domain open-close motion (10, 28, 29) makes crystallographic and biophysical characterisation of this interaction challenging. As a result, only short fragments of the CTD bound to the GTase have been resolved, e.g. only one CTD heptad was resolved in the mammalian GTase-CTD structure (Figure 1B) (32). However, much longer CTD peptides are required in order to elicit the stimulation of GTase activity, suggesting that a more extensive GTase-CTD interaction must occur (12). There are a number of positively charged regions in the mammalian CE GTase that have the potential to form additional pSer interaction sites (Figure 1C). In addition, the available GTase-CTD cocrystal structures provide no insights into the GTase allosteric activation mechanism, with the GTase conformations in the cocrystal structures being almost identical to their CTD-unbound equivalents (30–32). Nor do the current structural studies identify the differences between the pSer5 and pSer2 CTD interactions with the GTase that could explain why the activation effect is observed with the pSer5 CTD but not the pSer2 CTD (12). Therefore, there are a number of outstanding questions in the field: i) how does the CTD phosphorylation code affect CTD binding to the GTase?; ii) what additional GTase-CTD interactions are required for GTase activation; iii) how does pSer5 CTD binding to the GTase illicit GTase activity stimulation?

Computational techniques such as molecular dynamics (MD) simulations are well posed to answer these open questions and generate a more comprehensive characterisation of the GTase-CTD interaction (33). MD simulations are increasingly used in the characterisation of protein conformational dynamics and protein-peptide interactions, including energetics. Recent studies highlight the application of MD simulations to understand the conformational ensembles of protein systems, protein allostery and protein-peptide interactions (35–40). It must be mentioned that atomistic MD simulations are computationally expensive, typically limiting simulations to nanosecond-microsecond timescales (41). Since many events in protein systems occur on longer timescales, enhanced sampling techniques have been developed and successfully applied to overcome these limitations and sample longer-timescale processes (42, 43). Accelerated molecular dynamics (aMD) is one such technique that increases the conformational sampling of a system by reducing the depth of free-energy minima while maintaining the characteristics of the energy surface (44–48).

Here we carry out a large-scale computational study by performing both conventional MD (cMD) and accelerated MD simulations to assess the conformational dynamics of the human CE GTase and provide a systematic and detailed characterisation of its interaction with the CTD in different phosphorylation states. We identify two novel CTD interaction sites on the human CE GTase surface. These sites are predominantly conserved throughout animals and yeasts, indicating that the core features of the GTase-CTD interface have undergone considerably higher selection pressure than previously recognised. In addition, we propose that the GTase-CTD interaction is bidirectional and recognise the palindromic nature of the CTD.

## 3 Methods

### System preparation

The 3.0 Å resolution crystal structure of the human CE GTase (residues 229–565) (10) was used in simulations of the GTase systems. The systems prepared for MD simulations were constructed in PyMOL (49) based upon crystal structures of CE GTases available in the Protein Databank (PDB): i) the human CE *apo*-GTase (PDB ID: 3S24, Chain F) (10), ii) the mouse CE GTase in complex with one CTD heptad (PDB ID: 3RTX, Chains B and C) (32), iii) the *C. albicans* CE GTase in complex with ∼2.5 CTD heptads (PDB ID: 1P16, Chain B and D) (31), and iv) the *Paramecium bursaria Chlorella* virus 1 (PBCV-1) *holo*-GTase domains (PDB ID: 1CKM, Chain A) (28). All simulations were performed in the *apo* state, i.e. without ligands, in the presence or absence of the CTD. The human *apo* CE GTase crystal structure has seven GTase molecules in the asymmetric unit, varying in their conformational states between the ‘open’ and ‘closed’ conformation of the active site cleft. All simulations were started from the most open state. Supplementary Table 1 provides a summary of all the simulations presented in this work, with details of each system setup described below. All current crystal structures of the mammalian GTase miss portions of the *β*2-*α*D loop (residues 425–433). This was modelled with ModLoop using the MODELLER loop modelling procedure (50).

To simulate the CE GTase-CTD complex, the mouse GTase-CTD complex structure and human *apo*-GTase structure were aligned and the 1-heptad CTD fragment was superimposed onto the human CE GTase. To model the 4-heptad systems, the PEP-FOLD3.5 server (51) was used to generate the starting peptide structures of 3 additional heptads (21 residues). These 3 heptads were then fused onto the 1-heptad CTD resolved in the mouse GTase-CTD cocrystal structure, either onto its C- or N-terminus (Supplementary Table 1). Unique starting conformations were used for each replicate of the simulation by selecting different PEP-FOLD3.5-generated structures.

All simulations were prepared within the LEaP module in the AMBER16 suite (52) using the ff14SB force field (53), with phosphoserine modifications described by Homeyer *et al.* (54). All protein and peptide chains were capped with acetyl (ACE) and amino (NME) groups on the N- and C-ter respectively, and Reduce was used to protonate all residues in their standard protonation state at neutral pH (55). All CTD phosphoserines were modelled in the *−*2 charge state. Simulations of the 4-heptad, pSer5 CTD system were also performed with the pSer in the *−*1 charge state and show comparable qualitative behaviour, though with a weaker interaction and reduced stability of sites (data not shown). The protein was then placed in an octahedral box of TIP3P water molecules extending at least 15Å from the protein. The system was neutralised by balancing the charge with the appropriate number of Na^+^ or Cl*^−^* counter ions. Finally a combination of steepest descent and conjugate gradient energy minimisation was performed.

### Simulation setup and protocols

All standard simulations were performed using the pmemd.cuda module of AMBER16 (52). After energy minimisation, the system was heated from 100 K to 310 K over 25 ps, restraining the solute. Equilibration was performed for 200 ps with the solute restraints gradually removed. After 200 ps of equilibration, hydrogen mass repartitioning was performed and the step size was increased from 2 to 4 fs for the production runs (56). Berendsen barostat and thermostat were used to keep pressure and temperature constant (1 atmosphere and 310 K) during the simulations (57). The non-bonded interaction cutoff distance was set to 10.0Å and the SHAKE algorithm used to restrain hydrogen bond lengths (58). To reduce neutralising counterion clustering around the phosphate groups of the CTD, a 20Å distance restraint (*k* = 20.0 kcal/mol *·* A) was imposed between all sodium counterions and phosphorus atoms of the CTD phosphoserines. Three replicates of each production run were performed by randomly generating the starting velocities.

aMD runs were performed with the AMBER16 implementation using the ‘dual-boost’ protocol as described previously (46, 47, 52, 59, 60). Briefly, this applies a potential energy boost to all atoms and an additional dihedral boost to torsion angles. The mean potential and torsion energies of each system was calculated from the last 50 ns of each 200 ns cMD replicate. These were then used to calculate the aMD parameters (*E_P_, α_P_, E_D_, α_D_*) based upon the guidelines described by Pierce *et al.* (46).

### Data analysis

VMD and PyMOL were used to inspect and visualise the trajectories (49, 61). Analysis of the MD trajectories was performed primarily in the CPPTRAJ module of the AMBER16 suite to compute interatomic distances, solvent exposure, root-mean-square deviations (RMSDs) and root-mean-square fluctuations (RMSFs) (52). Interatomic distances between CTD residues and the GTase residues were computed using the closest CTD residue from any of its heptads. All trajectories were analysed by using frames saved every 40 ps. Electrostatic potentials were generated using the Adaptive Poisson-Boltzmann Solver (APBS) implemented in the PyMOL APBS tools (62). Normal mode analysis was performed using the ElNémo web server using the default settings (63). Multiple sequence alignments were performed in Jalview using the Clustal Omega algorithm with default settings (64, 65), selecting only reviewed protein sequences from the NT domain InterPro family (IPR001339).

The binding free energy analysis was performed using the Molecular Mechanics-Generalised Born Surface Area (MMGBSA) method using the MMPBSA.py package (66) and following the protocol described by Genheden *et al.* (67). The final snapshots of the aMD simulations were taken from the 3 replicates of each system. These snapshots were used as starting structures for 50 x 200 ps simulations. MMGBSA analysis, including per residue decomposition, was then performed using snapshots from these simulations with an 8 ps time step. *In silico* mutagenesis was performed on the final aMD snapshots followed by 50 ns of unrestrained equilibration. The MMGBSA protocol was then performed on the mutant structures as described above.

An additional search for potential CTD binding sites on the CE GTase was performed with the PIPER-FlexPepDock global protein-peptide docking server (68). Default settings were chosen for all conditions. The server does not accept non-standard residues, therefore, glutamates were used as phosphomimetics to replace the CTD phosphoserines.

Disorder prediction of the CE sequence was performed using the MetadisorderMD2 server (69).

### Expression and purification of recombinant proteins

The DNA sequences of the human CE GTase and each sequence variant were synthesised and subcloned into the PGEX6p1-C-His plasmid vector by Thermo Fisher GeneArt. The PGEX6p1-C-His vector contains an N-terminal HRV 3C cleavable tag and a C-terminal hexahistidine tag. These plasmids were then transformed into BL21 (DE3) *E. coli* and were cultured in 200 mL of Power Broth (Molecular Dimensions) at 37 °C until A600 was between 0.6–0.8. Protein expression was induced with 1mM IPTG overnight at 16 °C. Cells were pelleted and frozen before protein purification at *−*80 °C. The cells were lysed in 5 mL of lysis buffer (50 mM Tris-HCl, pH 7.5, 500 mM NaCl, 30 mM imidazole, 1 mM TCEP, 0.2% Tween and 5 units/ML Benzonase nuclease) and sonicated for 10 minutes with 10 second pulses. The GTase was purified with metal affinity chromatography, through a 1 mL HisTrap HP column (GE Healthcare) and eluting with 350 mM imidazole. The GST was cleaved with GST-tagged HRV 3C protease (PreScission Protease, GE Healthcare). The GST and protease was removed with glutathione sepharose resin. Further purification was performed with size exclusion chromatography on a Superdex 75 10/300 GL column (GE Healthcare), resolving in a buffer of 20 mM Tris HCl (pH 7.5), 200 mM NaCl and 1 mM TCEP. Aliquots were stored with 10% glycerol. Purity was assessed by SDS-PAGE and Coomassie Blue protein staining and all recombinant proteins were tested for basal GTase activity as described below (Supplementary Figure S7).

### CTD pull down assays

GTase-CTD peptide binding assays were performed as described by Ho *et al.* (12). 1 nmol of biotinylated 4-heptad CTD peptides (PeptideSynthetics) were incubated with 0.5 mg of Streptavidin-coupled magnetic Dynabeads M-280 (Invitrogen) in 300 *µ*L of buffer A (25mM Tris-HCl, pH 8, 50 mM NaCl, 1 mM TCEP, 5% glycerol and 0.03% Triton X-100) for 45 minutes at 4 °C. Next, the beads were magnet concentrated and washed three times with 0.5 mL of buffer A. 4 *µ*g of the purified GTase sample was then incubated with the beads in 50 *µ*L buffer B (Tris-HCl, pH 8, 53mM NaCl, 1 mM TCEP, 5% glycerol and 0.03% Triton X-100) for 45 minutes at 4 °C. After incubation, the solution was collected as the unbound fraction, the beads were washed three times with buffer A and the bound fraction was eluted with 50 *µ*L of SDS-PAGE loading buffer at 100 °C for 5 minutes. Fractions were concentrated and analysed with SDS-PAGE and Coomassie Blue staining. Bands were quantified in ImageJ and normalised relative to the wild-type CE GTase (residues 211–597) incubated with the Ser5 phosphorylated CTD peptide.

### Guanylyltransferase activity assays

Guanylyltransferase activity assays were performed as described by Ghosh *et al.* (32). 1 *µ*M of purified human CE GTase was incubated for 1 hour with CTD peptides of different concentrations (0, 2.5, 5, 10, 20, 40, 60, 80 and 100 *µ*M) in a buffer of 20 mM Tris HCl (pH 8.0) and 50 mM NaCl. After incubation, the guanylyltransferase activity assay was initiated by adding 2 *µ*L of the GTase-CTD mixture into a total volume of 20 *µ*L of assay buffer. The final activity assay buffer was composed of 0.1 *µ*M CE GTase, 20 mM Tris HCl (pH 8.0), 50 mM NaCl, 5 mM DTT, 0.2 *µ*M GTP (10% *α*^32^P, Perkin Elmer), 5 mM MgCl_2_ with varying concentrations of 4-heptad CTD peptide (0, 0.25, 0.5, 1, 2, 4, 6, 8 and 10 *µ*M). Reaction mixtures were incubated at 37 °C for 10 minutes and quenched with 1x loading buffer at 65 °C for 10 minutes. 15 *µ*L of each sample was run on an SDS-PAGE gel. The gels were fixed with 30% methanol and 5% acetic acid, stained with Coomassie Blue and exposed to a phosphorimaging plate for 1 hour. The plates were scanned using an Amersham Typhoon phosphorimager with the bands quantified in ImageJ and normalised relative to the basal wild-type CE GTase (residues 211–597) activity.

## 4 Results and Discussion

### The human CE GTase exhibits different conformational dynamics from the viral enzyme

To our knowledge the human Capping Enzyme guanylyltransferase domain (CE GTase) has not been simulated before. Therefore, our first aim was to assess the conformational dynamics of the protein. To characterise the conformational changes involving the GTase subdomains, the human CE GTase was simulated starting from the open state, without the CTD bound, and running 200 ns of cMD followed by 200 ns of aMD. In all replicates the structure enters the closed conformation and remains stable for the duration of the simulations (Supplementary Figure S1A). The NT and OB subdomains remain quasirigid, with RMSDs below 5 Å (Supplementary Figures S1B and S1C), similar to previous studies of GTase structures (28, 29, 70). As expected, these fluctuations are higher during the aMD simulations.

An important feature of GTase domains is the large-scale open-closed transition of the active site cleft, which is required for substrate binding, catalysis and product release (10, 28, 29). A previous computational study investigated the *Paramecium bursaria Chlorella* virus (PBCV-1) CE GTase and showed that the *apo* state can readily adopt the closed, open and hyperopen conformations (29). In contrast, in our simulations the *apo* human CE GTase samples the open and hyperopen states only briefly before becoming stabilised in the closed state (Figure 2A, Supplementary Figures S1A and S1D). For comparison, we also ran simulations of the PBCV-1 GTase, and focused our analysis on the interatomic domain distance. In contrast to the human CE GTase, the PBCV-1 GTase readily adopts the open and hyperopen states, fully consistent with the results of Swift *et al.* (29) (Figures 2A and 2B). This confirms that the two enzymes indeed exhibit strikingly different global dynamics.

**Figure 2.**
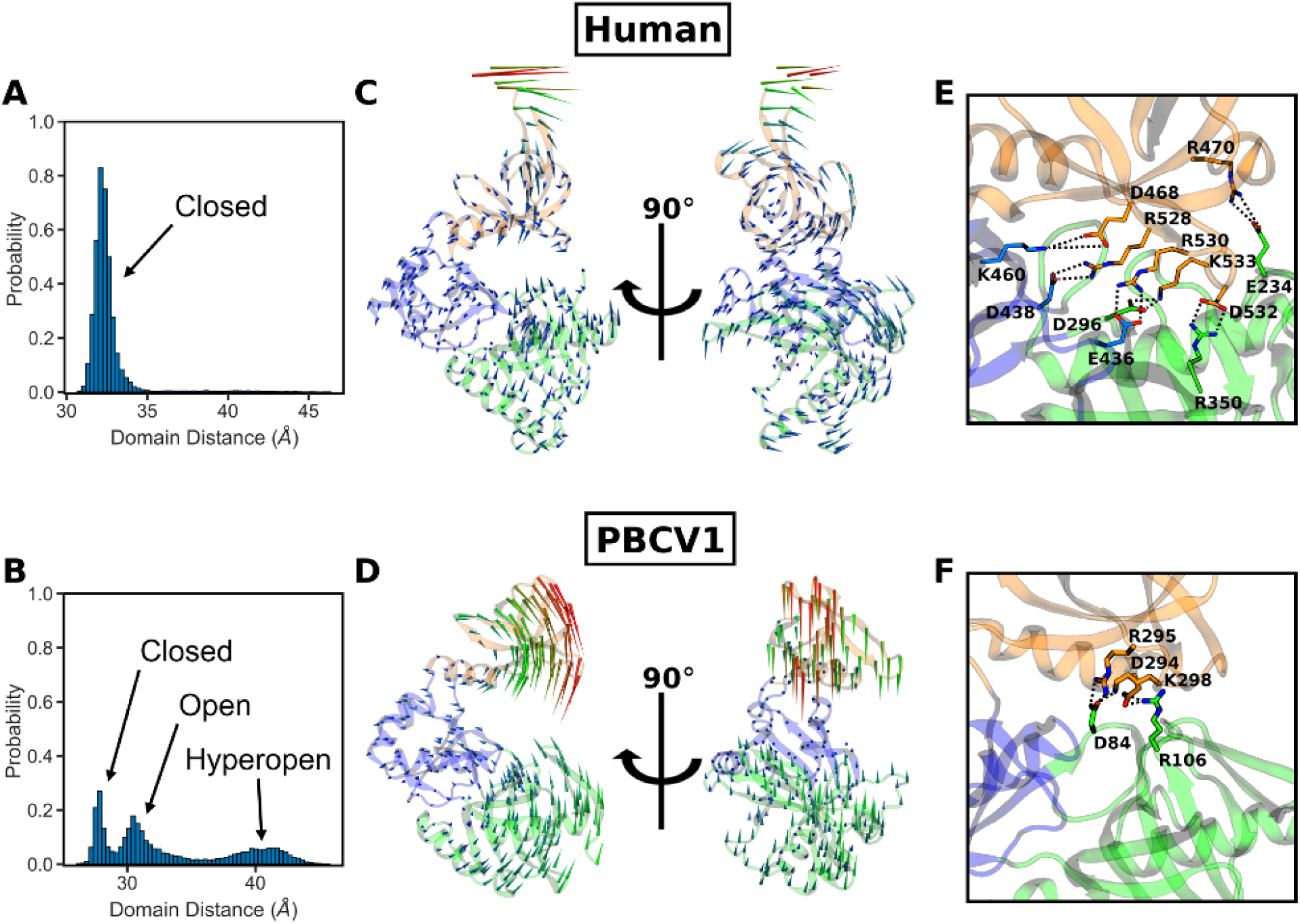
Conformational dynamics of the *apo* human CE GTase domain in comparison with the *apo* PBCV-1 CE GTase. Top panels (**A, C, E**) refer to the human GTase (10), and lower panels (**B, D, F**) refer to the PBCV-1 GTase (28). (**A-B**) Conformational distributions obtained from cMD simulations. The inter-domain distance was taken as the separation between the centres of mass of the NT and OB domains. Histograms were constructed using the data from all cMD replicates. (**C-D**) The results of the normal mode analysis, visualising conformational changes in the protein. The lowest frequency mode relating to the open-close motion is depicted as a porcupine plot, with the arrows representing the direction and amplitude of motion for each residue. (**E-F**) Salt bridges forming between the OB and NT domains of the GTase upon domain closure.

To further characterise these large-scale conformational changes Normal Mode Analysis (NMA) was performed on the human and PBCV-1 GTase structures (Figures 2C and 2D). The NMA results provide additional support to the above result, showing that for both structures the lowest frequency modes involve the domain opening and closing motion. However, this mode differs significantly between the two proteins. In the human CE GTase it is a rotation of the OB and NT domains relative to each other (Figure 2C). In contrast, for the PBCV-1 CE GTase the lowest frequency mode shows a straight open-close motion (Figure 2D). These differences in the global conformational dynamics are likely a result of the number of salt bridges that are able to form between the NT and OB domains (Figures 2E and 2F). In the human CE GTase, there is a complex network of salt bridges which hold the domains in the closed state, whereas in the PBCV-1 CE GTase no more than three salt bridges can be observed at any point during the simulations. Interestingly, a number of residues involved in salt bridge formation between the NT and OB domains in the human CE GTase—namely K460, D468, R528, R530, D532 and K533—have also been shown to be important residues for GTP and mRNA binding and mammalian CE GTase catalysis (10, 71, 72). However, the enzyme kinetics of the CE GTase have only been characterised in PBCV-1 and not the human CE GTase (73). The observed dramatic differences in the domain opening/closing dynamics between the two proteins suggest that the kinetics of the human enzyme will be significantly altered compared to that of the viral enzyme.

### The CTD forms an extensive interaction with the CE GTase, including two novel sites

The interaction between the CE GTase and the C-terminal domain of RNA Polymerase II (CTD) is essential for GTase activation and CE recruitment to the site of transcription (12, 15, 17). It has previously been shown that interactions with multiple heptads are required for GTase activation (12). As a starting point for understanding the interaction between the human CE GTase and the CTD, we initially carried out MD simulations of the GTase in the presence of 1 heptad of the CTD. These simulations were started from the CTD conformation and phosphorylation state observed in the mouse GTase-CTD cocrystal structure, which resolved only 1 heptad, phosphorylated at both the Ser2 and Ser5 positions (32). Our cMD results were consistent with the previous experimental data (Supplementary Figure S2): pSer5 remains bound to the positively charged pocket formed by R330, K331 and R386 (CDS1 site) in the conformation adopted in the crystal structure; in contrast, the pSer2 sidechain remains solvent exposed and does not form stable interactions with the protein. During the aMD simulations the CTD peptide samples much wider conformational space (Supplementary Figure S2). While the pSer5 interaction remains predominantly stable, the pSer2 residue changes conformation allowing it to also occasionally interact with the pSer5 pocket, CDS1. In addition, Tyr1 exhibits a greater extent of conformational flexibility, dissociating and rebinding to the tyrosine binding site (CDS-Y1).

Activation of the mammalian GTase strongly depends on the length of the CTD it interacts with, with the activation effect increasing 3-fold from 2 to 6 heptads (12). Activation is also dependent on the CTD being phosphorylated at the Ser5 position (12, 32). This indicates that the CTD forms an extensive interaction with the GTase that requires the binding of multiple CTD heptads. Currently there are no crystal structures of the mammalian CE GTase in complex with multiple CTD heptads. In order to systematically characterise the extensive interaction between the longer CTD fragments and the human CE GTase, we extended the length of the CTD peptide to 4 heptads by modelling 3 additional heptads onto the termini of the 1-heptad CTD fragment, which was resolved in the mouse crystal structure (32), in both directions. To investigate the effect of the CTD phosphorylation code on the GTase-CTD interaction, three phosphorylation states were simulated: unphosphorylated, Ser5 and Ser2 phosphorylated (Supplementary Table S1). In each phosphorylation state the CTD peptide was extended in both the N- and C-ter directions in separate simulation systems, yielding 6 different systems, to identify interaction sites that might occur at different sides of the known CTD interaction sites (CDS1 and CDS-Y1) (Supplementary Figure S3). Three replicates were performed using different CTD starting conformations to ensure that the interactions formed were reproducible and not biased by the initial CTD conformation (Supplementary Figure S3).

Analysis of the 4-heptad pSer5 CTD simulations (Systems 6 and 7) provided valuable insights into the GTase-CTD interaction (Figure 3). The previously reported CDS1 site remains occupied in all replicates (Figure 3B). The CDS-Y1 interaction remains stable for the duration of the cMD simulations but becomes destabilised, dissociating and rebinding, during the aMD simulations (Figure 3D). In addition to CDS1 and CDS-Y1, our simulations identify two novel CDS sites—named CDS2 and CDS-Y2—that were not observed in the mouse crystal structure of the complex (Figure 3 and Movie S1) (32).

**Figure 3.**
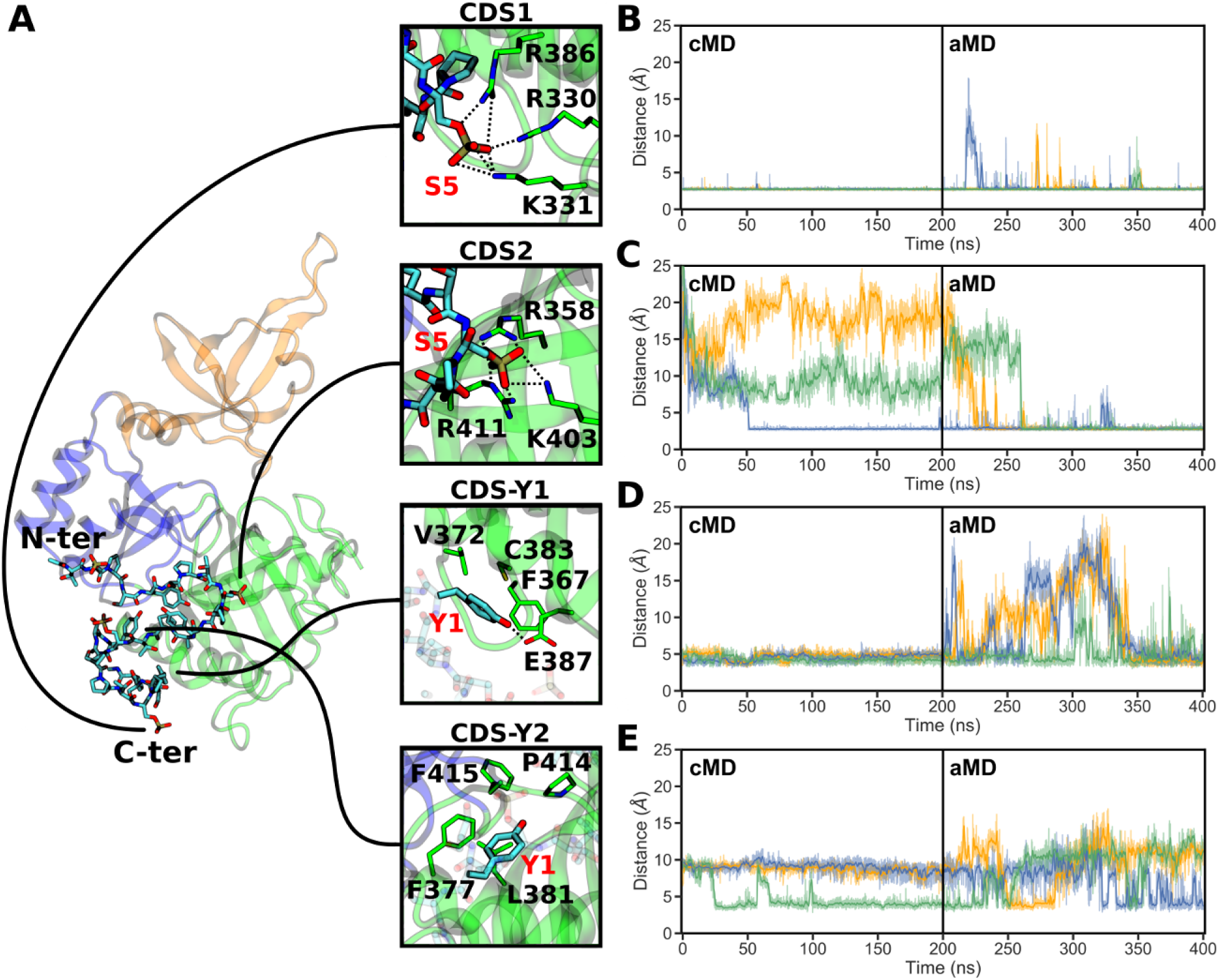
CTD interaction sites observed during the simulations of the GTase with the 4-heptad pSer5 CTD extended in the N-ter direction (System 6). (**A**) Locations on the GTase (left) and residues involved in the four CTD interaction sites (panels on the right) reproducibly observed in these simulations. The snapshot shows a representative CTD binding conformation and the positions of the N- and C-terminal heptads are indicated. (**B-E**) Time-evolution of the minimum distances showing the occupation of each CDS by the respective CTD group (pSer5 or Tyr1) over the duration of the cMD and aMD simulations. Distances obtained in three replicates are shown in orange (replicate 1), blue (replicate 2) and green (replicate 3). The occupation of each site was described by taking representative sidechain minimum distances as follows: (**B**) CDS1, R330 sidechain nitrogens to the pSer5 phosphate oxygens, (**C**) CDS2, R411 sidechain nitrogens to the pSer5 phosphate oxygens, (**D**) CDS-Y1, V372 C*γ* atoms to the Tyr1 ring, and (**E**) CDS-Y2, L381 sidechain to the Tyr1 ring. Distances from the CTD groups to other residues in the respective pockets (e.g. R386 and K331 in CDS1) show comparable behaviour.

The first novel CDS site, CDS2, is a pSer5 interaction site composed of sidechains R358, K403 and R411 that form multiple electrostatic interactions with the phosphate group of pSer5 of the CTD (Figures 3A, 3C and Movie S2). The CDS2 site is located within a positively charged patch on *β*7, *β*8 and loop *α*C-*β*8. This interaction is very stable, remaining occupied once pSer5 binds to the site, and is observed in all replicates of System 6 (Figure 3C). In the no-CTD state, the basic residues that constitute the CDS2 site are predominantly solvent exposed and are involved in transient interactions with surrounding negatively charged groups, including D349, D402, E406 and E432. No large-scale conformational changes occur upon pSer5 binding to CDS2, ruling out an induced fit mechanism.

The second novel CDS site identified by simulations is a tyrosine pocket, CDS-Y2. This accommodates the Tyr1 residue of the CTD, through hydrophobic interactions of the tyrosine ring with L381 at the centre of the pocket and F377, P414 and F416 in the vicinity (Figures 3A, 3E and Movie S3). This pocket is partially occupied by Tyr1 in all replicates, however, it represents a transient interaction and was easily destabilised during the aMD simulations (Figure 3E). The CDS-Y2 residues are located on helix *α*C and loop *β*8-*α*D. They are semi-buried within the NT-Hinge interface, reducing their solvent exposure, and interact with a number of adjacent hydrophobic residues that form part of a hydrophobic region that includes W293, Y362, I384, F416 and T445. Although there is no large-scale conformational change associated with Tyr1 binding to this site, many of these residues interact directly with the residues involved in GTP binding. Therefore Tyr1 binding to this site might have an effect on GTP binding or coordination.

Due to their electrostatic nature, the pSer5-CDS interactions (CDS1 and CDS2 sites) remain stable once formed (Figures 3B and 3C). During aMD simulations, some individual CDS1 interactions are occasionally broken, however pSer5 remains bound to this region. Upon CDS2 binding, this interaction remains stable with only minor fluctuations. In contrast, the Tyr1 interactions are considerably less stable (Figures 3D and 3E). CDS-Y1 remains occupied by Tyr1 for the duration of the cMD, however all replicates show Tyr1 dissociation and rebinding during the aMD stage. This is also observed with the CDS-Y2 pocket, which again represents a transient interaction, despite being occasionally observed in all replicates.

A further inspection of the previous cocrystal structure of the mouse GTase-CTD complex provides a rationale to explain why the newly identified CTD interaction sites, i.e. CDS2 and CDS-Y2, were not observed in that structure (32). The asymmetric unit of the structure forms a homodimer between two GTase domains, which was considered an artefact of crystallisation (Supplementary Figure S4). This homodimer interface forms extensive contacts on the NT domain and the hinge, occluding the CDS2 and CDS-Y2 sites, close to the bound CTD heptad. As a result, the dimer interface obstructs the CDS2 and CDS-Y2 sites, preventing CTD binding to this region. We expect that future structural studies of the mammalian GTase will confirm CTD binding to these novel sites.

An important feature of our simulations is that although the novel CDS2 and CDS-Y2 interactions are observed reproducibly in all replicates (Figures 3C and 3E), these interactions can occur on different heptads between the replicates (Figure 4). The CDS1 and CDS2 sites can be occupied either by adjacent CTD heptads or heptads can be looped out, with non-neighbouring heptads occupying CDS1 and CDS2. This provides evidence of the ‘looping out’ mechanism suggested in previous studies, which showed that the CE must interact with multiple heptads but that these do not need to be adjacent in sequence (31). The simulations also show that the order of the CDS interactions can vary. This can be seen, for example, in replicate 2 where heptad 4 dissociates from CDS-Y1 and is replaced by heptad 3, switching the order of CDS1 and CDS-Y1 (Figure 4). Both conformations are stable and this change does not destabilise other CDS interactions. Therefore, CDS sites can be occupied in different heptad orders as well as heptads being looped out. During GTase recruitment the CTD is not uniformly Ser5 phosphorylated (25), and so the looping out mechanism we observe is consistent with the hypothesis that unphosphorylated CTD heptads are looped out during GTase recruitment to enable the CTD to bind to all the CDS sites (31).

**Figure 4.**
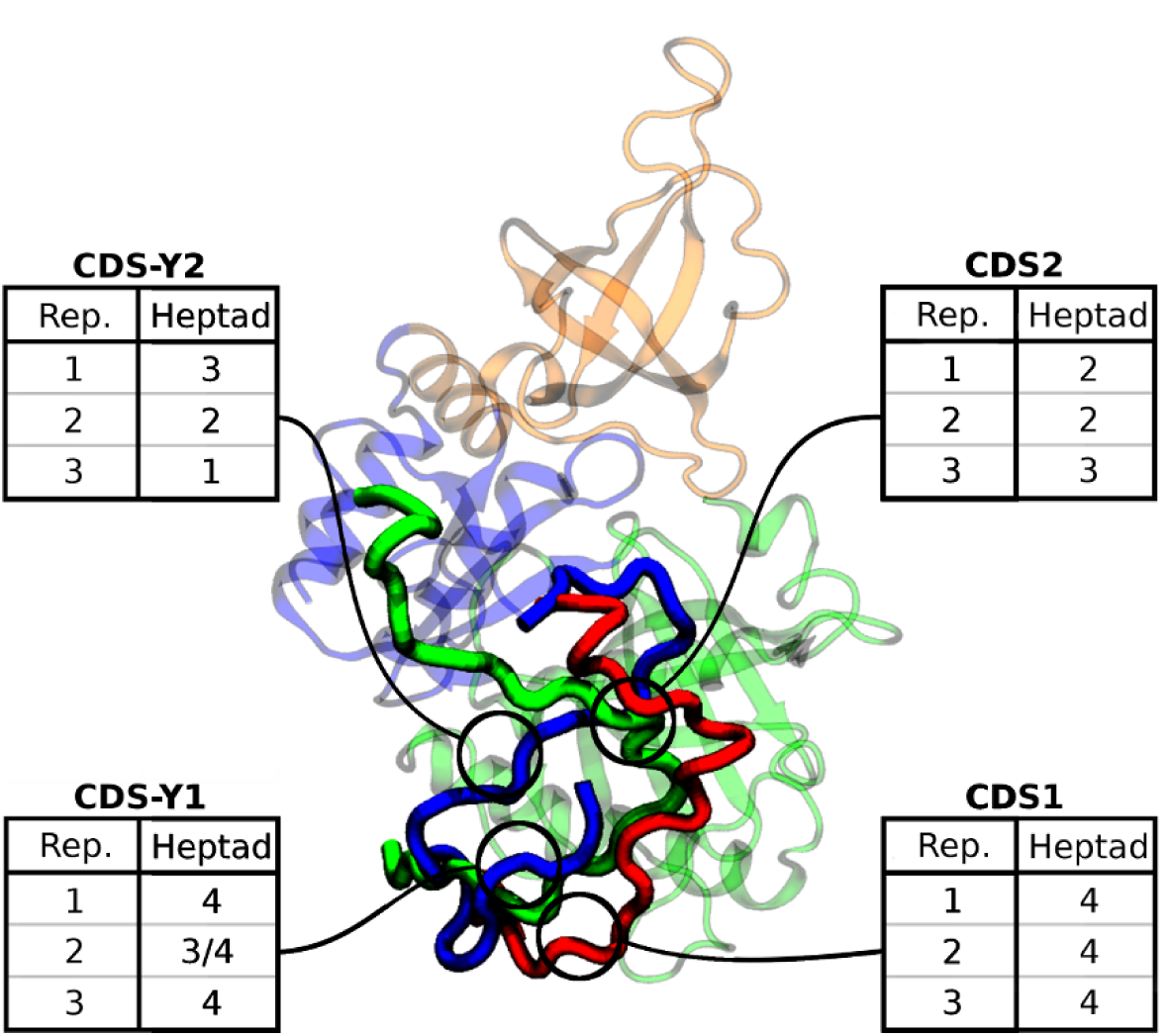
CTD peptide binding conformations and looping out mechanism. The CTD conformations are taken from the final aMD snapshots of the 4-heptad (N-ter extended), pSer5 CTD system (System 6), with the CTD in replicate 1 displayed in red, replicate 2 in blue, and replicate 3 in green. The four CDS sites are labelled by circles. The boxes indicate which heptad predominantly interacts with the respective site in each simulation replicate. CTD heptads are numbered from the N-terminus (1) to the C-terminus (4).

### Phosphoserine interaction sites are critical for CTD binding to the GTase

Our simulations revealed two additional CTD interaction sites on the human CE GTase surface. However, the contribution and importance of each CDS site to the CTD binding to the GTase remained unclear. MMGBSA is a computational technique that can be used to predict the binding free energies between binding partners, including protein-peptide complexes (see Methods for details) (66, 67, 74, 75). In order to obtain a detailed quantitative characterisation of the GTase-CTD interaction, MMGBSA calculations were performed to assess the binding affinities and the contributions of individual residues. The MMGBSA analysis identified the main GTase residues that contribute to CTD binding (Figure 5). Results for the 4-heptad pSer5 simulations extended in the N-ter direction (System 6) are shown in Figure 5A. As expected, the core residues comprising the pSer5 interaction sites—residues R330 and R386 of CDS1 and R358, K403 and R411 of CDS2—make the largest contributions to the GTase-CTD interaction. Notably, arginines make the most significant contribution to the binding free energy, whereas the flexibility of the CDS lysine sidechains and their position on the loops in the CE GTase make them more likely to dissociate from CTD interactions. This can be seen in CDS1 where R330 and R386 make the largest contributions to the CTD binding affinity, whereas K331 makes a relatively small contribution. Likewise, in CDS2, R358 and R411 make the largest contributions to the binding affinity, whereas K403 makes a smaller contribution because of its location on a loop. R392 is included as a CDS2 residue, however it forms a strong interaction with pSer5 only in one replicate, whereas in the other two replicates it forms a stable salt bridge with E406 in the NT domain; this explains a large standard deviation for this residue. No other residues on the CE GTase make significant contributions to the binding affinity, confirming the central role of CDS1 and CDS2 sites in the GTase-CTD interaction.

**Figure 5.**
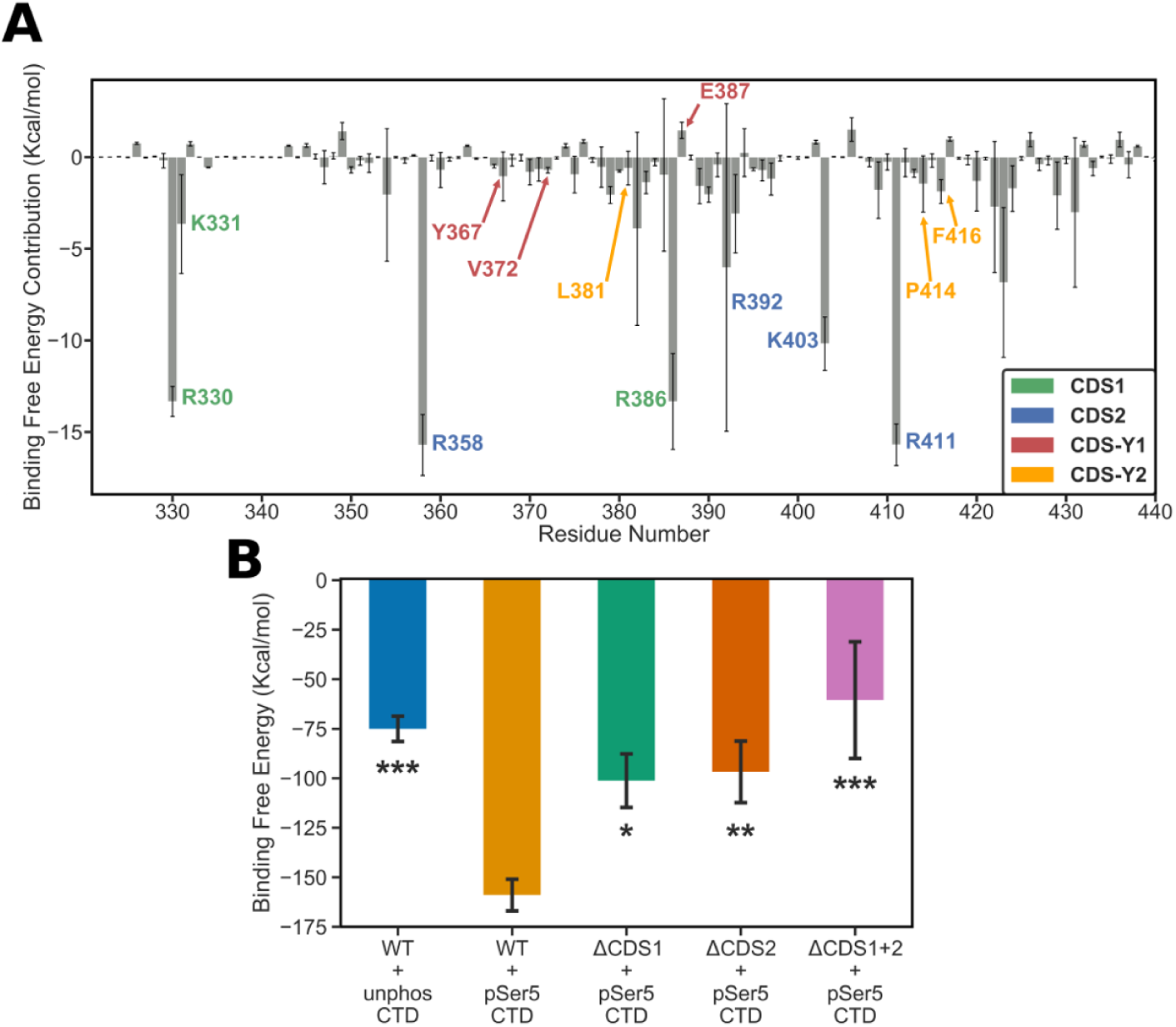
Quantitative analysis of the CTD binding affinity to the GTase. MMGBSA calculations (see details in the Methods) were performed on three replicates of the simulations containing the 4-heptad pSer5 CTD extended in the N-ter direction (System 6). (**A**) Per-residue decomposition analysis of the binding free energy. The key GTase residues that contribute to CTD binding are labelled and coloured according to the CDS site they belong to (see the plot legend box). The residues making significant contributions (below −2.1 kcal/mol) are all confined to the region between residues 320 and 440. (**B**) Comparison of the binding free energy between the wild-type GTase and the three mutants, where the positively charged residues of CDS1, CDS2 and both sites were mutated to alanine. The result for the wild-type GTase with unphosphorylated CTD is also shown for comparison. Error bars denote one standard deviation. ANOVA followed by post-hoc Tukey tests were performed to calculate statistical significance between the mutants and the wildtype GTase + pSer5 CTD condition. * indicates that the differences are significant at P*<*0.05, ** indicates that the differences are significant at P*<*0.01, and *** indicates that the differences are significant at P*<*0.001.

In contrast to the pSer5 sites, the Tyr1 sites make a minor contribution to the binding affinity, with none of the CDS-Y1 or CDS-Y2 residues contributing more than *−*5 kcal/mol (Figure 5A). This is consistent with their transient nature seen in the aMD distance analysis (Figures 3D and 3E). Despite this, mutagenesis of Tyr1 to alanine has been previously shown to significantly decrease GTase binding and activation (76). This suggests that Tyr1 has an essential but more subtle role in GTase recruitment and activation.

As pSer5 interactions were found to dominate the GTase-CTD interaction, *in silico* mutagenesis was performed to further assess the importance of each site and to guide biochemical experiments (Figure 5B). We constructed mutant systems in which CDS1 (R330, K331 and R386) and CDS2 (R358, K403 and R411) residues were mutated to alanine in the final frames of the aMD simulations of System 6—yielding Systems 13, 14 and 15 (Supplementary Table 1). Each system was reequilibrated for 50 ns and then MMGBSA analysis was performed (Figure 5B). An additional system containing the dephosphorylated CTD (System 12) was simulated to provide an important reference. It must be noted that although MMGBSA results are extremely useful to compare relative binding free energy values for different systems, the absolute binding free energy values calculated must be taken with caution (67). The results show that the two pSer5 CDS sites (CDS1 and CDS2) make major contributions to the binding affinity of the CTD. When either pocket is mutated (ΔCDS1 or ΔCDS2) the binding free energy is reduced by about a third. When residues in both CDS1 and CDS2 pockets are mutated to alanine, the binding free energy is reduced further and is approximately equal to that of the unphosphorylated CTD. These results suggest that, although the Tyr1 interactions may have have an auxiliary role in GTase recruitment and activation, pSer5 CDS interactions form the basis of GTase binding to the CTD.

### pSer2 CTD can bind to CDS1 and CDS2 adopting different conformations from pSer5 CTD

In order to characterise the differences in the GTase-CTD interaction as the CTD code is changed, we simulated the pSer2 CTD (Systems 8 and 9) and compared its conformational dynamics and interactions to that of the pSer5 CTD (Systems 6 and 7). pSer2 is known to bind to the GTase with comparable affinity to pSer5 but does not illicit the GTase activation (12). Previous literature suggests that the Ser2 phosphorylated CTD displays non-competitive binding with Ser5 phosphorylated CTD, therefore, the two states are expected to bind to different locations on the CE GTase surface (12). Our MD results suggest that the pSer2 CTD also readily binds to the same sites, CDS1 and CDS2 (Figure 6 and Supplementary Figure S5). During the simulations, the CDS1 pocket is quickly occupied by pSer2 due to its close starting proximity (Figure 6B and Supplementary Figure S5A). In addition, in one of the three replicates extended in the N-ter direction and in two of the three replicates extended in the C-ter direction the pSer2 sidechain occupies the CDS2 pocket (Figure 6A, 6C and Supplementary Figure S5B). Once pSer2 is bound, the respective sites remain occupied for the duration of the simulations. These dynamics are similar to the pSer5 CTD. This indicates that the CDS1 and CDS2 pockets are not specific to pSer5, and that both pSer2 and pSer5 groups can bind to them. Importantly, the conformation the pSer2 CTD adopts when binding to the CDS2 site is different from that of the pSer5 CTD (Figure 6A). As the pSer2 residue is adjacent to Tyr1, it reduces the Tyr1 interactions with the hydrophobic CDS-Y1 and CDS-Y2 pockets (Figures 6D and 6E, Supplementary Figures S5C and S5D). Tyr1 interactions have been implicated in CE recruitment and activation by the CTD (76). Therefore, this difference in binding mode may explain why pSer2 CTD can bind to the human CE GTase but does not stimulate GTase activity (12).

**Figure 6.**
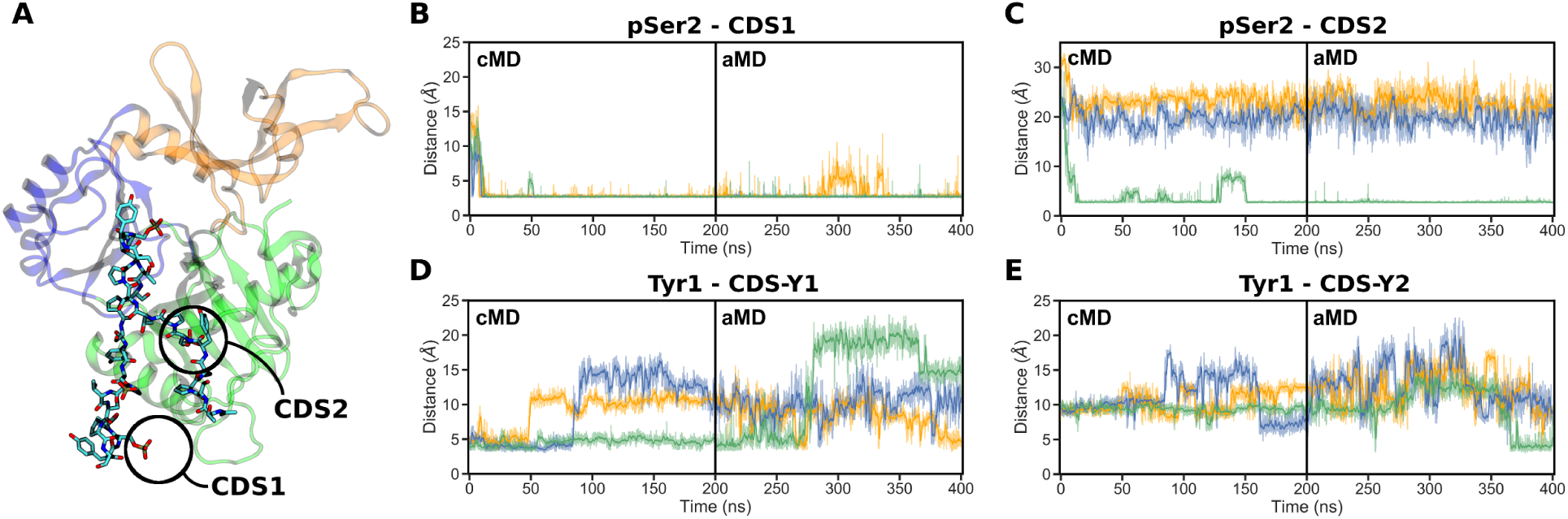
CTD interaction sites observed during the simulations of the GTase with the 4-heptad pSer2 CTD extended in the N-ter direction. (**A**) The final snapshot of replicate 3 showing the CTD bound to both CDS1 and CDS2 interaction sites on the GTase. (**B-E**) Time-evolution of the minimum distances showing the occupation of each CDS by the respective pSer2 CTD group over the duration of the cMD and aMD simulations. Distances obtained in three replicates are shown in orange (replicate 1), blue (replicate 2) and green (replicate 3). The occupation of each site was described by taking representative sidechain minimum distances as follows: (**B**) CDS1, R330 sidechain nitrogens to the pSer2 phosphate oxygens, (**C**) CDS2, R411 sidechain nitrogens to the pSer2 phosphate oxygens, (**D**) CDS-Y1, V372 C atoms to the Tyr1 ring, (**E**) CDS-Y2, L381 sidechain to the Tyr1 ring.

### Disordered flanking domains contain positively charged regions suitable for phosphorylated CTD binding

Our simulations provide a detailed understanding of the CTD interactions with the GTase within a distance of around 3 heptads from the site reported in the mouse GTase-CTD crystal structure (32). However, they do not account for interactions that could occur in more distant regions of the human CE, such as the OB domain or within the disordered regions at the N- and C-terminal flanks of the GTase, which were not resolved in any of the crystal structures (10, 32). A previous crystal structure of the *S. pombe* CE GTase (Pce1) displayed a Spt5 CTD docking site in the OB-fold domain (30). Given that the full-length CTD is 52 heptads in humans, it is not excluded that some fragments of the CTD also interact with other regions of the human CE, even when bound to the CDS1 and CDS2 sites (19).

The task of exploring potential binding sites of a full-length CTD on the CE GTase is unfeasible for atomistic MD. Therefore, to explore potential binding sites in alternative regions of the GTase, global peptide docking was performed using the PIPER-FlexPepDock server. Although global protein:peptide docking is challenging and often inaccurate, these techniques can give an indication of the regions a peptide can bind to and the possible conformations it can adopt. In particular, the results for the 2-heptad CTD docking to the GTase show that the pSer5 CTD peptides are localised in the NT domain in the region covering CDS1 and CDS2 (Supplementary Figure S6B). On the other hand, the docking results for the pSer2 CTD offer a less clear picture although they still show models docked to CDS2 (Supplementary Figure S6C). These observations provide additional support to our MD findings.

So far we have focussed on the CTD binding to the CE GTase domain. However, CTD binding to other regions of the CE has not been fully explored. Biochemical assays have previously shown that the phosphorylated CTD does not interact with the TPase domain of the human CE, but the contribution of the two disordered regions that flank the GTase domain has not been examined previously (12). Unfortunately, due to the length and the disordered nature of these regions, they could not be accurately modelled or simulated by MD simulations. Inspection of the human CE sequence reveals that both the disordered TPase-GTase linker and the disordered region at the C-terminus of the GTase contain large numbers of positively charged residues that are found in several clusters, which could form positively charged sites similar to CDS1 and CDS2 (Supplementary Figure S6D). Therefore, we expect that the phosphorylated CTD can interact with these regions in addition to the sites in the GTase domain, enhancing the CE recruitment to the CTD. The C-terminal flanking region has previously been shown to be essential for the recruitment of the CE to Nck1 to enable cytoplasmic capping, further suggesting that this region plays an important role in recruitment of the CE to both the site of cotranscriptional and cytoplasmic capping (77).

### Biochemical assays validate the role of the phosphoserine interaction sites for GTase recruitment

Our computational analyses provide a comprehensive picture of the GTase-CTD interaction offering a number of findings that can be tested biochemically. In particular, our results predict that: i) pSer interactions with the CE GTase form the basis of the binding affinity, with CDS1 and CDS2 sites making major contributions to CTD binding affinity, ii) mutating out both CDS1 and CDS2 reduces CTD binding affinity to a level comparable to the unphosphorylated CTD, iii) CDS1 and CDS2 are non-specific and can also bind the Ser2 phosphorylated CTD, and iv) the disordered regions flanking the GTase domain contain positively charged residues that are likely to contribute to CTD binding. In order to validate these predictions, we expressed and purified the recombinant human GTase proteins and tested the affinity of the 4-heptad CTD peptides (Supplementary Figure 7). A total of eight recombinant proteins were prepared: the core human CE GTase (229–569) wild-type, ΔCDS1 (R330A/K331A/R386A), ΔCDS2 (R358A/K403A/R411A) and ΔCDS1+2 (R330A/K331A/R386A/R358A/K403A/R411A). In addition, WT and mutant proteins were expressed and purified containing both disordered domains that flank the GTase (211–597).

First, we performed pull-down assays on all recombinant proteins, using 4 heptad CTD peptides that were either unphosphorylated, Ser5 or Ser2 phosphorylated on all heptads (Figures 7A-7E). The WT GTase (211–597) binds to the 4 heptad pSer5 CTD with an affinity comparable with previous literature (Figure 7A) (12). We then compared the WT CE with the core human GTase domain (229–569) and the GTase with the disordered flanking domains (211–597) (Figures 7A and 7B). In agreement with our prediction that these disordered regions might be important for GTase recruitment, the protein containing the flanking regions exhibits a significantly increased CTD binding. These interactions are not pSer5 or pSer2 specific, enhancing binding for both the pSer5 and pSer2 CTD peptides. As both the CTD and these flanking regions are disordered, these additional interactions possibly represent the formation of a ‘fuzzy’ complex where the CTD interacts at well-ordered sites on the GTase surface (CDS sites) in addition to forming interactions with the disordered flanking regions (78, 79). In this case the role of the flanking interactions is to increase GTase-CTD binding required for CE recruitment and GTase activation.

**Figure 7.**
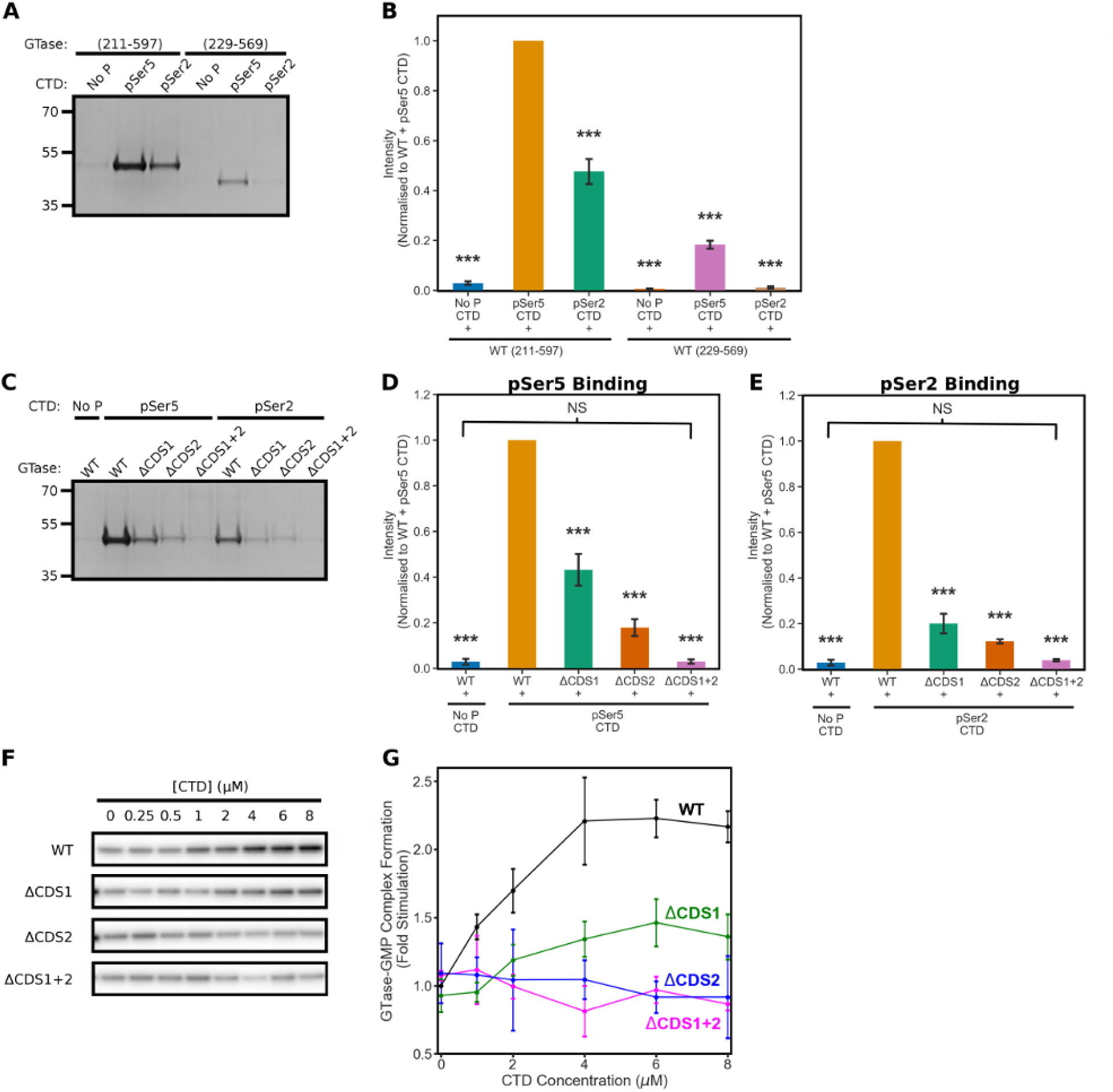
Biochemical assays characterising GTase binding to the CTD and GTase activity stimulation. (**A-E**) Recombinant human CE GTase was incubated with biotinylated peptides of 4 CTD heptads—in either their unphosphorylated, Ser5- or Ser2-phosphorylated states—bound to streptavidin-coupled Dynabeads. The level of GTase binding to the CTD peptides was assessed by SDS-PAGE stained with Coomassie Blue. (**A-B**) Comparison of the constructs with and without the disordered flanks and with the 4-heptad CTD peptides (unphos, pSer2 or pSer5). (**C-E**) Com-parison of the CTD binding affinity to the GTase (211-597) mutants for the unphosphorylated, pSer5 and pSer2 CTD peptides. (**F-G**) Guanylyltransferase activity assay of the GTase (211-597) mutants with increasing CTD concentration. Band quantification of the assays—shown in (B), (D), (E) and (G)—was performed in triplicate and the mean was plotted, normalising to the WT + pSer5 CTD condition. Error bars denote one standard deviation. ANOVA followed by post-hoc Tukey tests were performed to calculate statistical significance compared to the wild-type GTase (211-597) + pSer5 CTD condition. *** indicates that the differences are significant at P *<* 0.001.

We then sought to validate the pSer5 CTD interactions observed on the GTase domain. Comparison of the binding affinity of the CDS mutations shows results consistent with our computational predictions, with the CDS1 and CDS2 sites both contributing significantly to pSer5 CTD binding (Figures 7C and 7D, Supplementary Figure 8). When both of these interaction sites are removed, GTase binding to pSer5 CTD is at a comparable level with the WT GTase binding to the unphosphorylated CTD. This indicates that there are no other pSer5 interaction sites on the GTase.

In further agreement with our computational results, pSer2 CTD peptide binding also significantly decreases when CDS1 or CDS2 residues are mutated, with the same trend observed for the pSer5 CTD peptide (Figures 7C and 7E). This result contrasts with previous literature that showed that the pSer2 CTD binds to the human CE GTase non-competitively with the pSer5 CTD (12). Our results also show that pSer2 CTD has a lower binding affinity than pSer5 (Figures 7A and 7B), in contrast with previous experimental data that showed the pSer2 and pSer5 CTD peptides binding to the CE GTase with comparable affinity (12).

These results are reproducibly observed in the core GTase assays (229–569), however, without the disordered flanking regions the low binding affinity makes it difficult to quantify and distinguish the mutants (Supplementary Figure 6).

### The novel phosphoserine pocket, CDS2, is essential for GTase activation

Upon binding to the CE GTase, the Ser5 phosphorylated CTD has been shown to stimulate the first step of GTase catalysis (12). All crystal structures of the CE GTase-CTD interaction show that the CTD binds outside of the active site, on the NT domain, therefore this must involve an allosteric mechanism of activation. The nature of such an allosteric activation, its mechanism and importance in the regulation of mRNA capping remain unclear (26, 27). To assess whether the novel CDS2 site is involved in the CE GTase activation by the CTD, we performed GTase activity assays on the CDS mutant recombinant GTase proteins (Figures 7F and 7G). The activity assay quantifies the first stage of GTase activity by measuring the level of *α*^32^P labelled guanosine monophosphate covalently bound to the GTase active site. Incubation of the WT GTase (211–297) with the 4 heptad pSer5 CTD increases the GTase activity by 2.2-fold, consistent with previous literature (12). Removal of the CDS1 site reduces the GTase activation effect, however it still elicits activation to 1.4 fold the basal level. In contrast, mutagenesis of the CDS2 site completely inhibits GTase activity stimulation, suggesting that it has an essential role in the allosteric activation of the GTase. Mutagenesis of both the CDS1 and CDS2 sites replicates the inhibition observed in the ΔCDS2 mutant.

To explore the molecular details of allosteric activation, we compared the conformational dynamics of the GTase in its pSer5 CTD-bound and no-CTD states (Supplementary Figure S9 and S10). Comparison of the aMD simulations of these two systems shows no global changes in the GTase secondary structure (Supplementary Figures S9A and S9B), in agreement with previous crystal structures (31, 32). In addition, the aMD simulations show no significant changes in the dynamics of the GTase (Supplementary Figure S10). However, upon binding of the CTD to the CDS2 pocket, some local conformational changes do occur. In particular, the conformational dynamics of loop *β*9-*α*D, which is predominantly unstructured in the no-CTD state, favours an *α*-helical arrangement upon pSer5 CTD binding (Supplementary Figures S9C-S9F). This loop lies adjacent to the Mg^2+^ binding site and the GTP binding pocket and therefore this structural rearrangement might alter Mg^2+^ and GTP binding affinity or coordination, increasing the rate of the first step of GTase catalysis. However, a full assessment of the role of CTD binding on GTase activation is beyond the scope of this work, requiring comprehensive analysis of CTD binding at the GTP-bound and intermediate stages of GTase activation. This will be the focus of the subsequent work.

### The GTase-CTD interaction sites are predominantly conserved between animals and yeasts

After identification of novel CTD interaction sites in the simulations, we checked these in the available crystal structures of the GTase-CTD complex in other eukaryotic species (*S. pombe* and *C. albicans*) (30, 31). Previous research has concluded that different taxa have evolved distinct CDS binding sites on the GTase surface to recruit the CE to the site of transcription (30, 32): although all current GTase-CTD cocrystal structures show that the CTD interacts with the NT domain of the CE GTase, their conformations and interaction sites differ significantly (30–32). Despite this, *pombe* and *C. albicans* GTase-CTD cocrystal structures share a number of conserved features, including a CDS site (CDS1) composed of residues on helix *α*C and loop *β*7-*α*C and a Tyr1 interaction site in the same location between helix *α*C and strand *β*8. Apart from these similarities, they contain additional pSer5 interaction sites that are not conserved between the two species or seen in the mouse GTase-CTD cocrystal structure. When comparing these yeast GTase-CTD interactions with the mouse GTase-CTD cocrystal structure there are no conserved interactions between them, although the CTD sites are always in nearby regions of the NT subdomain (32).

Surprisingly, when comparing the novel CDS sites observed in our simulations of the human CE GTase with that in the *C. albicans* cocrystal structure, we find a number of similarities. Both the novel CDS2 and CDS-Y2 sites are also observed at the same positions in the *C. albicans* cocrystal structure (Figures 3A and 8A). In addition, CDS2 residues have been shown to be essential for CTD binding to the *S. cerevisiae* CE GTase (80). Sequence analysis comparing animal and yeast species shows that the core residues of both CDS2 and CDS-Y2 are functionally conserved across animals and yeasts (Figure 8B). For the CDS2 site this functional conservation is not immediately apparent because, although R358 is highly conserved throughout animals and yeasts, K403 and R411 are not conserved in yeasts. However, in yeasts both are substituted with nearby positively charged residues that are highly conserved: K403 is substituted with a positively charged residue on helix *α*C (K178 in *C. albicans*) and R411 is substituted for a lysine two residues away on the same side of strand *β*8 (K193 in *C. albicans*). These residues are not conserved in *S. pombe* and the CDS2 site is not observed in its GTase-CTD cocrystal structure (30), suggesting divergent evolution in this branch of yeasts.

**Figure 8.**
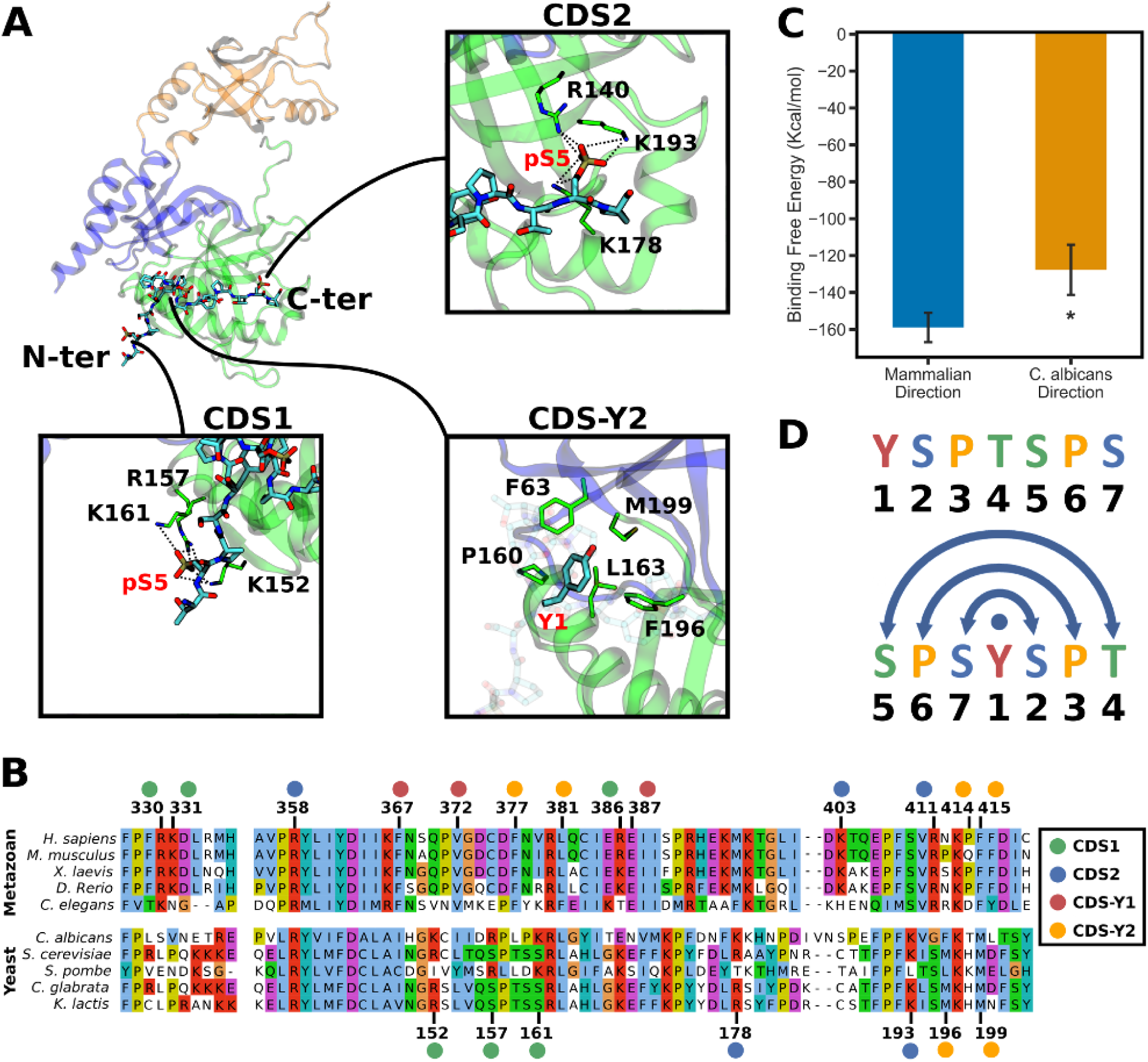
Conservation analysis of the CTD interaction sites on the GTase, comparison to the *C. albicans* GTase-CTD cocrystal structure, and the palindromic nature of the CTD. (**A**) CTD interaction sites observed in the *C. albicans* GTase-CTD cocrystal structure (PDB ID: 1P16) (31). The positions of the N- and C-ter heptads of the CTD are indicated. The residue numbers are from the *C. albicans* GTase. (**B**) Multiple sequence alignment of animal and yeast CE GTase sequences in the CDS regions. Numbers above indicate the human CE residue numbers and below are the *C. albicans* residue numbers. The CDS sites are indicated by coloured circles above/below the residue numbers. (**C**) MMGBSA analysis comparing the CTD binding affinity to the human GTase between the “mammalian” orientation (as in Figure 5; System 6) and the “yeast” orientation (from the *C. albicans* crystal structure extended to four heptads; System 11; see details in the Methods). (**D**) One CTD heptad displayed in the conventional way (above) and then shifted and centred at Tyr1 (below) to illustrate the palindromic nature of the repeating sequence.

The CDS-Y2 pocket identified in our simulations is also conserved throughout animals and yeasts. The central leucine residue (L381) is highly conserved between the two. The additional hydrophobic residues that comprise the pocket are highly conserved in animals and yeasts but the specific residues in this pocket vary between the two taxa. In animals, this pocket is composed of F377, P414 and F415, in contrast to F63, F196 and M199 in yeasts.

The CDS1 pocket, although its precise location in loop *β*5-*β*6 and helix *α*C is not conserved between animals and yeasts, is in the same region of the GTase across animals and yeasts. The residues in the CDS1 pocket are highly conserved in animals, however, the CDS1 residues are poorly conserved throughout yeasts. Despite this, most species of yeast contain positively charged residues in either loop *β*5-*β*6, where the mammalian CDS1 residues are located, or on loop *β*7-*α*C and helix *α*C, the same location as in *C. albicans* and *S. pombe*. Interestingly, R386 is highly conserved as a positively charged residue throughout both animals and yeasts. As this residue is only around 7Å from the CTD pSer5 sidechain in the *S. pombe* cocrystal structure, it is likely that this residue also contributes to CTD binding in yeasts. This lack of conservation of the CDS1 site is unsurprising because the majority of the CDS1 residues are located on flexible loops where the exact position of the positively charged residues is unlikely to affect efficient GTase recruitment by the CTD. The CDS-Y1 hydrophobic pocket, in contrast to the CDS-Y2 pocket, is highly conserved in animals, however, does not appear to be conserved in yeasts and it is not observed in either of the yeast GTase-CTD cocrystal structures (30, 31). Therefore, although many of the central features of the GTase-CTD interaction are conserved in both animals and yeasts, there are some features that distinguish them.

### The palindromic nature of the CTD code allows bidirectional binding of the CTD

One striking difference between the conformations seen in our simulations and those observed in the *C. albicans* CE GTase-CTD cocrystal structure is that the CTD peptide is oriented in opposite directions (Figures 3A and 8A). In our simulations CDS1 is occupied by a pSer5 of the C-terminal heptad of the CTD and CDS2 by a pSer5 of an N-terminal heptad. In contrast, the *C. albicans* cocrystal structure shows CDS1 occupied by the N-terminal heptad and CDS2 by the C-terminal heptad. This raised the question of whether this is a characteristic feature that is distinct between mammals and yeasts or if the CTD can bind in both directions.

To assess the stability and affinity of CTD binding to the human CE GTase in the alternative orientation, the *C. albicans* CTD conformation was superimposed onto the human CE GTase, extended to 4 heptads and simulations were performed as described above (Systems 10 and 11; Supplementary Figure S11). The CTD conformation remains stable for the duration of the simulations, with the same characteristics as observed in our previous simulations: the pSer5 pockets (CDS1 and CDS2) form strong interactions and the CDS-Y2 site is more transient (Supplementary Figure 7). Thus the pSer5 CTD can bind to the same sites in both orientations, although the CTD with the “mammalian” orientation has a higher relative binding affinity than the CTD bound in the “yeast” orientation (Systems 6 and 11; Figure 8C).

We suggest that such bidirectional CTD binding to the same interaction sites is enabled because the CTD heptad motif is almost completely palindromic (Figure 8D), and therefore the positions of the CTD residues remain the same in both directions. To our knowledge this feature of the CTD sequence and structure has not been previously discussed. To see this, the canonical heptad motif must be viewed starting with Ser5 and placing Tyr1 at the centre. The GTase-CTD interaction mostly involves the CTD sidechains, such as the pSer and Tyr1 interactions, but not the backbone, and therefore the chirality of the peptide backbone is unlikely to affect the GTase-CTD binding affinity. This also agrees with our finding that the CDS pockets can be occupied in different heptad orders (Figure 3B). As a result, the *C. albicans* CTD conformation can be superimposed onto the human GTase with few minor steric clashes, and this conformation remains stable for the duration of the simulation, despite the fact that it is in the “opposite orientation”. Bidirectionality of peptide binding has been observed for other protein-peptide interactions, including the WW domain, MHC class II, SH3 domain and O-GlcNAcase (81–84). In particular, the WW domain in Pin1 binds to CTD phosphopeptides in an opposite direction to other examples of WW domain protein-peptide interactions (81). Bidirectional peptide binding has been suggested to have implications for binding specificity, as changes to the peptide sequence or phosphorylation pattern are likely to introduce steric constraints that may prevent binding in a particular orientation (82).

We hypothesise that the palindromic nature of the CTD contributes to its function in binding to such a wide variety of partners during transcription. It may allow the CTD to have specific interactions, such as interactions with the specific CTD kinases, while also being able to recruit factors such as the CE, where the CTD can bind in a variety of conformations. The palindromic nature of the CTD also has implications for how the code could be read, with the pSer5 and pSer2 being in distinct locations along the palindrome. This could, in part, explain why a major transition in phosphorylation occurs between Ser5 and Ser2 rather than Ser2 and Ser7, which are in the same relative position within the palindrome (20, 22, 25, 85).

## 5 Conclusions

Recruitment of the Capping Enzyme (CE) by the carboxyl-terminal domain of RNA Polymerase II (CTD) is an essential stage of mRNA capping, localising the CE to the site of transcription and stimulating the activity of its guanylyltransferase (GTase) domain (15–17). Despite a number of studies of the GTase-CTD interface, fundamental questions remain about the molecular details of this interaction (12, 27, 30–33, 76, 80). We have carried out an extensive and systematic study of the GTase-CTD interaction using molecular dynamics (MD) simulations, in which we varied the phosphorylation code, length and orientation of the CTD. We have subsequently confirmed the main computational predictions by performing a series of biochemical assays.

Through this approach we have identified several distinct characteristics of the GTase-CTD interaction. Most notably, we have identified two novel interaction sites on the human CE GTase surface (CDS2 and CDS-Y2). In addition to this we have shown that the disordered flanks of the GTase contribute significantly to CTD binding. Structural and sequence analysis between animals and yeasts reveals that the novel GTase-CTD interaction sites are highly conserved, leading us to conclude that the GTase-CTD interaction sites have undergone considerably higher selection pressure than previously considered. The binding free energy analysis and binding assays have demonstrated that the phosphoserine interactions are the main contributors to the GTase-CTD interaction. Subsequent activity assays have shown that the novel CDS2 site is essential for GTase activation, revealing a previously missing link for understanding the molecular mechanism of GTase activation by the CTD.

This work has also characterised how the GTase-CTD interaction depends on the CTD phosphorylation code. Our simulations and biochemical assays both show that these interactions are not pSer5 CTD specific and are also essential for pSer2 CTD binding. We conclude that the occupation of the pSer interaction sites does not confer allosteric activation alone, instead the distinct conformations the pSer5 CTD peptides adopt when bound at these sites determines whether the GTase becomes activated. Overall, this work moves forward our current understanding of the GTase-CTD interaction, from one of static interaction sites obtained from crystal structures to a more complex picture of transient interactions and structural ensembles, where the CDS sites are occupied in different orders and directions.

Finally, this work sheds light on the structural features of the CTD. Our simulations clearly demonstrate the CTD looping out mechanism first described by Fabrega *et al.* (31). Moreover, we show that the order of heptad binding to the CDS sites can change more drastically, leading to the identification of the GTase-CTD interaction bidirectionality. We conclude that this bidirectionality is the result of the CTD motif being palindromic. The palindromic nature of the CTD has not been explored previously but it is likely to have implications for how the CTD is written and read, affecting a number of stages of gene regulation.

## 6 Acknowledgements

We are grateful to Professor Tom Owen-Hughes and Dr Subbu Sundaramoorthy for kindly providing the PGEX6p1-C-His plasmid vector and technical advice on protein expression and purification. We would also like to thank Professor Ulrich Zachariae for critical discussions.

## References

1. Ramanathan, A., Robb, G. B. & Chan, S.-H. mRNA capping: biological functions and applications. Nucleic acids research 44, 7511–26 (2016).

2. Galloway, A. & Cowling, V. H. mRNA cap regulation in mammalian cell function and fate. Biochimica et Biophysica Acta (BBA) - Gene Regulatory Mechanisms 1862, 270–279 (2019).

3. Izaurralde, E. et al. A nuclear cap binding protein complex involved in pre-mRNA splicing. Cell 78, 657–668 (1994).

4. Ohno, M., Sakamoto, H. & Shimura, Y. Preferential excision of the 5’ proximal intron from mRNA precursors with two introns as mediated by the cap structure. Proceedings of the National Academy of Sciences of the United States of America 84, 5187–91 (1987).

5. Andersen, P. R. et al. The human cap-binding complex is functionally connected to the nuclear RNA exosome. Nature structural & molecular biology 20, 1367–76 (2013).

6. Jiao, X., Chang, J. H., Kilic, T., Tong, L. & Kiledjian, M. A mammalian pre-mRNA 5’ end capping quality control mechanism and an unexpected link of capping to pre-mRNA processing. Molecular cell 50, 104–15 (2013).

7. Cheng, H. et al. Human mRNA Export Machinery Recruited to the 5 End of mRNA. Cell 127, 1389–1400 (2006).

8. Choe, J. et al. eIF4AIII enhances translation of nuclear cap-binding complex-bound mRNAs by promoting disruption of secondary structures in 5’UTR. Proceedings of the National Academy of Sciences of the United States of America 111, E4577–86 (2014).

9. Changela, A., Ho, C., Martins, A., Shuman, S. & Mondragón, A. Structure and mechanism of the RNA triphosphatase component of mammalian mRNA capping enzyme. The EMBO Journal 20, 2575–2586 (2001).

10. Chu, C. et al. Structure of the guanylyltransferase domain of human mRNA capping enzyme. Proceedings of the National Academy of Sciences of the United States of America 108, 10104– 8 (2011).

11. Ho, C. K. et al. The guanylyltransferase domain of mammalian mRNA capping enzyme binds to the phosphorylated carboxyl-terminal domain of RNA polymerase II. The Journal of biological chemistry 273, 9577–85 (1998).

12. Ho, C. K. & Shuman, S. Distinct roles for CTD Ser-2 and Ser-5 phosphorylation in the recruitment and allosteric activation of mammalian mRNA capping enzyme. Molecular cell 3, 405–11 (1999).

13. Varshney, D. et al. Molecular basis of RNA guanine-7 methyltransferase (RNMT) activation by RAM. Nucleic acids research 44, 10423–10436 (2016).

14. Gonatopoulos-Pournatzis, T., Dunn, S., Bounds, R. & Cowling, V. H. RAM/Fam103a1 is required for mRNA cap methylation. Molecular cell 44, 585–96 (2011).

15. Yue, Z. et al. Mammalian capping enzyme complements mutant Saccharomyces cerevisiae lacking mRNA guanylyltransferase and selectively binds the elongating form of RNA polymerase II. Proc. Natl. Acad. Sci. USA 94, 12898–12903 (1997).

16. Moteki, S. & Price, D. Functional Coupling of Capping and Transcription of mRNA. Molecular Cell 10, 599–609 (2002).

17. McCracken, S. et al. 5’-Capping enzymes are targeted to pre-mRNA by binding to the phosphorylated carboxy-terminal domain of RNA polymerase II. Genes & development 11, 3306– 3318 (1997).

18. Cho, E.-J., Takagi, T., Moore, C. R. & Buratowski, S. mRNA capping enzyme is recruited to the transcription complex by phosphorylation of the RNA polymerase II carboxy-terminal domain. Genes & Development 11, 3319–3326 (1997).

19. Jeronimo, C., Collin, P. & Robert, F. The RNA Polymerase II CTD: The Increasing Complexity of a Low-Complexity Protein Domain. Journal of molecular biology 428, 2607–2622 (2016).

20. Harlen, K. M. & Churchman, L. S. The code and beyond: transcription regulation by the RNA polymerase II carboxy-terminal domain. Nature Reviews Molecular Cell Biology 18, 263–273 (2017).

21. McCracken, S. et al. The C-terminal domain of RNA polymerase II couples mRNA processing to transcription. Nature 385, 357–360 (1997).

22. Schüller, R. et al. Heptad-Specific Phosphorylation of RNA Polymerase II CTD. Molecular Cell 61, 305–314 (2016).

23. Komarnitsky, P., Cho, E. J. & Buratowski, S. Different phosphorylated forms of RNA polymerase II and associated mRNA processing factors during transcription. Genes & development 14, 2452–60 (2000).

24. Cho, E. J., Kobor, M. S., Kim, M., Greenblatt, J. & Buratowski, S. Opposing effects of Ctk1 kinase and Fcp1 phosphatase at Ser 2 of the RNA polymerase II C-terminal domain. Genes & development 15, 3319–29 (2001).

25. Suh, H. et al. Direct Analysis of Phosphorylation Sites on the Rpb1 C-Terminal Domain of RNA Polymerase II. Molecular cell 61, 297–304 (2016).

26. Noe Gonzalez, M., Sato, S., Tomomori-Sato, C., Conaway, J. W. & Conaway, R. C. CTD-dependent and -independent mechanisms govern co-transcriptional capping of Pol II transcripts. Nature Communications 9, 3392 (2018).

27. Schwer, B. & Shuman, S. Deciphering the RNA Polymerase II CTD Code in Fission Yeast. Molecular Cell 43, 311–318 (2011).

28. Håkansson, K., Doherty, A. J., Shuman, S. & Wigley, D. B. X-Ray Crystallography Reveals a Large Conformational Change during Guanyl Transfer by mRNA Capping Enzymes. Cell 89, 545–553 (1997).

29. Swift, R. V. & McCammon, J. A. Substrate Induced Population Shifts and Stochastic Gating in the PBCV-1 mRNA Capping Enzyme. Journal of the American Chemical Society 131, 5126–5133 (2009).

30. Doamekpor, S. K., Sanchez, A. M., Schwer, B., Shuman, S. & Lima, C. D. How an mRNA capping enzyme reads distinct RNA polymerase II and Spt5 CTD phosphorylation codes. Genes & development 28, 1323–36 (2014).

31. Fabrega, C., Shen, V., Shuman, S. & Lima, C. D. Structure of an mRNA capping enzyme bound to the phosphorylated carboxy-terminal domain of RNA polymerase II. Molecular Cell 11, 1549–1561 (2003).

32. Ghosh, A., Shuman, S. & Lima, C. D. Structural Insights to How Mammalian Capping Enzyme Reads the CTD Code. Molecular Cell 43, 299–310 (2011).

33. Burley, S. K. & Sonenberg, N. Gimme Phospho-Serine Five! Capping Enzyme Guanylyltransferase Recognition of the RNA Polymerase II CTD. Molecular Cell 43, 163–165 (2011).

34. Jasnovidova, O. & Stefl, R. The CTD code of RNA polymerase II: a structural view. Wiley Interdisciplinary Reviews: RNA 4, 1–16 (2013).

35. Hospital, A., Goñi, J. R., Orozco, M. & Gelpí, J. L. Molecular dynamics simulations: advances and applications. Advances and applications in bioinformatics and chemistry : AABC 8, 37–47 (2015).

36. . Huggins, D. J. et al. Biomolecular simulations: From dynamics and mechanisms to computational assays of biological activity. Wiley Interdisciplinary Reviews: Computational Molecular Science 9, e1393 (2019).

37. Mou, L. et al. Microsecond molecular dynamics simulations provide insight into the ATP-competitive inhibitor-induced allosteric protection of Akt kinase phosphorylation. Chemical Biology & Drug Design 89, 723–731 (2017).

38. Hertig, S., Latorraca, N. R. & Dror, R. O. Revealing Atomic-Level Mechanisms of Protein Allostery with Molecular Dynamics Simulations. PLOS Computational Biology 12, e1004746 (2016).

39. Ahuja, L. G., Aoto, P. C., Kornev, A. P., Veglia, G. & Taylor, S. S. Dynamic allostery-based molecular workings of kinase:peptide complexes. Proceedings of the National Academy of Sciences of the United States of America 116, 15052–15061 (2019).

40. Paul, F. et al. Protein-peptide association kinetics beyond the seconds timescale from atomistic simulations. Nature Communications 8, 1095 (2017).

41. Dror, R. O., Dirks, R. M., Grossman, J. P., Xu, H. & Shaw, D. E. Biomolecular Simulation: A Computational Microscope for Molecular Biology. Annu. Rev. Biophys 41, 429–52 (2012).

42. Bernardi, R. C., Melo, M. C. R. & Schulten, K. Enhanced sampling techniques in molecular dynamics simulations of biological systems. Biochimica et biophysica acta 1850, 872–877 (2015).

43. Miao, Y. & McCammon, J. A. Unconstrained enhanced sampling for free energy calculations of biomolecules: a review. Molecular Simulation 42, 1046–1055 (2016).

44. Hamelberg, D., Mongan, J. & McCammon, J. A. Accelerated molecular dynamics: A promising and efficient simulation method for biomolecules. The Journal of Chemical Physics 120, 11919–11929 (2004).

45. Bucher, D., Grant, B. J., Markwick, P. R. & McCammon, J. A. Accessing a hidden conformation of the maltose binding protein using accelerated molecular dynamics. PLoS computational biology 7, e1002034 (2011).

46. Pierce, L. C., Salomon-Ferrer, R., de Oliveira, C. A. F., McCammon, J. A. & Walker, R. C. Routine Access to Millisecond Time Scale Events with Accelerated Molecular Dynamics. Journal of Chemical Theory and Computation 8, 2997–3002 (2012).

47. Bueren-Calabuig, J. A., Bage, M. G., Cowling, V. H. & Pisliakov, A. V. Mechanism of allosteric activation of human mRNA cap methyltransferase (RNMT) by RAM: insights from accelerated molecular dynamics simulations. Nucleic Acids Research 47, 8675–8692 (2019).

48. Moffett, A. S., Bender, K. W., Huber, S. C. & Shukla, D. Molecular dynamics simulations reveal the conformational dynamics of Arabidopsis thaliana BRI1 and BAK1 receptor-like kinases. The Journal of biological chemistry 292, 12643–12652 (2017).

49. The PyMOL Molecular Graphics System, Version 1.9, Schrödinger, LLC.

50. Fiser, A. & Sali, A. ModLoop: automated modeling of loops in protein structures. Bioinformatics 19, 2500–2501 (2003).

51. Lamiable, A. et al. PEP-FOLD3: faster de novo structure prediction for linear peptides in solution and in complex. Nucleic Acids Research 44, W449–W454 (2016).

52. Case, D. A., Betz, R. M., Cerutti, D. S., Cheatham, T. E., Darden, T. A., Duke, R. E., Giese, S. J., Gohlke, H. et al. AMBER16 (2016).

53. Maier, J. A. et al. ff14SB: Improving the Accuracy of Protein Side Chain and Backbone Parameters from ff99SB. Journal of Chemical Theory and Computation 11, 3696–3713 (2015).

54. Homeyer, N., Horn, A. H. C., Lanig, H. & Sticht, H. AMBER force-field parameters for phosphorylated amino acids in different protonation states: phosphoserine, phosphothreonine, phosphotyrosine, and phosphohistidine. Journal of Molecular Modelling 12, 281–289 (2006).

55. Word, J., Lovell, S. C., Richardson, J. S. & Richardson, D. C. Asparagine and glutamine: using hydrogen atom contacts in the choice of side-chain amide orientation. Journal of Molecular Biology 285, 1735–1747 (1999).

56. Hopkins, C. W., Le Grand, S., Walker, R. C. & Roitberg, A. E. Long-Time-Step Molecular Dynamics through Hydrogen Mass Repartitioning. Journal of Chemical Theory and Computation 11, 1864–1874 (2015).

57. Berendsen, H. J. C., Postma, J. P. M., van Gunsteren, W. F., DiNola, A. & Haak, J. R. Molecular dynamics with coupling to an external bath. The Journal of Chemical Physics 81, 3684– 3690 (1984).

58. Miyamoto, S. & Kollman, P. A. Settle: An analytical version of the SHAKE and RATTLE algorithm for rigid water models. Journal of Computational Chemistry 13, 952–962 (1992).

59. Hamelberg, D., de Oliveira, C. A. F. & McCammon, J. A. Sampling of slow diffusive conformational transitions with accelerated molecular dynamics. The Journal of Chemical Physics 127, 155102 (2007).

60. de Oliveira, C. A. F., Hamelberg, D. & McCammon, J. A. Coupling Accelerated Molecular Dynamics Methods with Thermodynamic Integration Simulations. Journal of chemical theory and computation 4, 1516–1525 (2008).

61. Humphrey, W., Dalke, A. & Schulten, K. VMD: Visual molecular dynamics. Journal of Molecular Graphics 14, 33–38 (1996).

62. Baker, N. A., Sept, D., Joseph, S., Holst, M. J. & McCammon, J. A. Electrostatics of nanosystems: application to microtubules and the ribosome. Proceedings of the National Academy of Sciences of the United States of America 98, 10037–41 (2001).

63. Suhre, K. & Sanejouand, Y.-H. ElNemo: a normal mode web server for protein movement analysis and the generation of templates for molecular replacement. Nucleic Acids Research 32, W610–W614 (2004).

64. Waterhouse, A. M., Procter, J. B., Martin, D. M. A., Clamp, M. & Barton, G. J. Jalview Version 2-a multiple sequence alignment editor and analysis workbench. Bioinformatics 25, 1189–1191 (2009).

65. Sievers, F. et al. Fast, scalable generation of high-quality protein multiple sequence alignments using Clustal Omega. Molecular systems biology 7, 539 (2011).

66. Miller, B. R. et al. MMPBSA.py : An Efficient Program for End-State Free Energy Calculations. Journal of Chemical Theory and Computation 8, 3314–3321 (2012).

67. Genheden, S. & Ryde, U. The MM/PBSA and MM/GBSA methods to estimate ligand-binding affinities. Expert opinion on drug discovery 10, 449–61 (2015).

68. Alam, N. et al. High-resolution global peptide-protein docking using fragments-based PIPER-FlexPepDock. PLOS Computational Biology 13, e1005905 (2017).

69. Kozlowski, L. P. & Bujnicki, J. M. MetaDisorder: a meta-server for the prediction of intrinsic disorder in proteins. BMC Bioinformatics 13, 111 (2012).

70. Swift, R. V. & McCammon, J. A. Catalytically requisite conformational dynamics in the mRNA-capping enzyme probed by targeted molecular dynamics. Biochemistry 47, 4102– 4111 (2008).

71. Wen, Y., Yue, Z. & Shatkin, A. J. Mammalian capping enzyme binds RNA and uses protein tyrosine phosphatase mechanism. Proc. Natl. Acad. Sci. USA 95, 12226–12231 (1998).

72. Sawaya, R. & Shuman, S. Mutational analysis of the guanylyltransferase component of mammalian mRNA capping enzyme. Biochemistry 42, 8240–8249 (2003).

73. Soulière, M. F., Perreault, J.-P. & Bisaillon, M. Kinetic and thermodynamic characterization of the RNA guanylyltransferase reaction. Biochemistry 47, 3863–74 (2008).

74. Hou, T., Wang, J., Li, Y. & Wang, W. Assessing the Performance of the MM/PBSA and MM/GBSA Methods. 1. The Accuracy of Binding Free Energy Calculations Based on Molecular Dynamics Simulations. Journal of Chemical Information and Modeling 51, 69–82 (2011).

75. Weng, G. et al. Assessing the performance of MM/PBSA and MM/GBSA methods. 9. Prediction reliability of binding affinities and binding poses for proteinpeptide complexes. Physical Chemistry Chemical Physics 21, 10135–10145 (2019).

76. Pei, Y., Hausmann, S., Ho, C. K., Schwer, B. & Shuman, S. The length, phosphorylation state, and primary structure of the RNA polymerase II carboxyl-terminal domain dictate interactions with mRNA capping enzymes. The Journal of biological chemistry 276, 28075–82 (2001).

77. Mukherjee, C., Bakthavachalu, B. & Schoenberg, D. R. The Cytoplasmic Capping Complex Assembles on Adapter Protein Nck1 Bound to the Proline-Rich C-Terminus of Mammalian Capping Enzyme. PLoS Biology 12, e1001933 (2014).

78. Tompa, P. & Fuxreiter, M. Fuzzy complexes: polymorphism and structural disorder in proteinprotein interactions. Trends in Biochemical Sciences 33, 2–8 (2008).

79. Wright, P. E. & Dyson, H. J. Intrinsically disordered proteins in cellular signalling and regulation. Nature Reviews Molecular Cell Biology 16, 18–29 (2015).

80. Bharati, A. P. et al. The mRNA capping enzyme of Saccharomyces cerevisiae has dual specificity to interact with CTD of RNA Polymerase II. Scientific Reports 6, 31294 (2016).

81. Verdecia, M. A., Bowman, M. E., Lu, K. P., Hunter, T. & Noel, J. P. Structural basis for phosphoserine-proline recognition by group IV WW domains. Nature Structural Biology 7, 639–643 (2000).

82. Kuriyan, J. & Cowburn, D. Modular Peptide Recognition Domains in Eukaryotic Signalling. Annual Review of Biophysics and Biomolecular Structure 26, 259–288 (1997).

83. Günther, S. et al. Bidirectional binding of invariant chain peptides to an MHC class II molecule. Proceedings of the National Academy of Sciences of the United States of America 107, 22219–24 (2010).

84. Li, B., Li, H., Hu, C.-W. & Jiang, J. Structural insights into the substrate binding adaptability and specificity of human O-GlcNAcase. Nature Communications 8, 666 (2017).

85. Heidemann, M., Hintermair, C., Voß, K. & Eick, D. Dynamic phosphorylation patterns of RNA polymerase II CTD during transcription (2013).

