## Supplementary Data for "Identification of novel RNA Polymerase II CTD interaction sites on the mRNA Capping Enzyme"

- 1 Supplementary Table
- 11 Supplementary Figures
- 3 Supplementary Movies
- Representative snapshots (in a PDB format) for each simulation system

**Table S1.** Summary of the molecular dynamics simulations performed in this work

| | CE GTase | CTD | cMD | aMD |
|---|---|---|---|---|
| 1 | human | - | 3x 200 ns | 3x 200 ns |
| 2 | PBCV-1 | - | 3x 200 ns | - |
| 3 | human | 1 heptad, pSer5 and pSer2 | 3x 200 ns | 3x 200 ns |
| 4 | human | 4 heptads, unphos, extended from N-ter | 3x 200 ns | 3x 200 ns |
| 5 | human | 4 heptads, unphos, extended from C-ter | 3x 200 ns | 3x 200 ns |
| 6 | human | 4 heptads, pSer5, extended from N-ter | 3x 200 ns | 3x 200 ns |
| 7 | human | 4 heptads, pSer5, extended from C-ter | 3x 200 ns | 3x 200 ns |
| 8 | human | 4 heptads, pSer2, extended from N-ter | 3x 200 ns | 3x 200 ns |
| 9 | human | 4 heptads, pSer2, extended from C-ter | 3x 200 ns | 3x 200 ns |
| 10 | human | ~2.5 heptads, pSer5, in *C. albicans* conform. | 3x 200 ns | 3x 200 ns |
| 11 | human | 4 heptads, pSer5, from *C. albicans* conform. | 3x 50 ns | - |
| 12 | human | System 6 final conform., dephos. | 3x 50 ns | - |
| 13 | human ΔCDS1 | System 6 final conform., pSer5 | 3x 50 ns | - |
| 14 | human ΔCDS2 | System 6 final conform., pSer5 | 3x 50 ns | - |
| 15 | human ΔCDS1/CDS2 | System 6 final conform., pSer5 | 3x 50 ns | - |
| | | Total: | 6.75 $\mu$s | 5.40 $\mu$s |

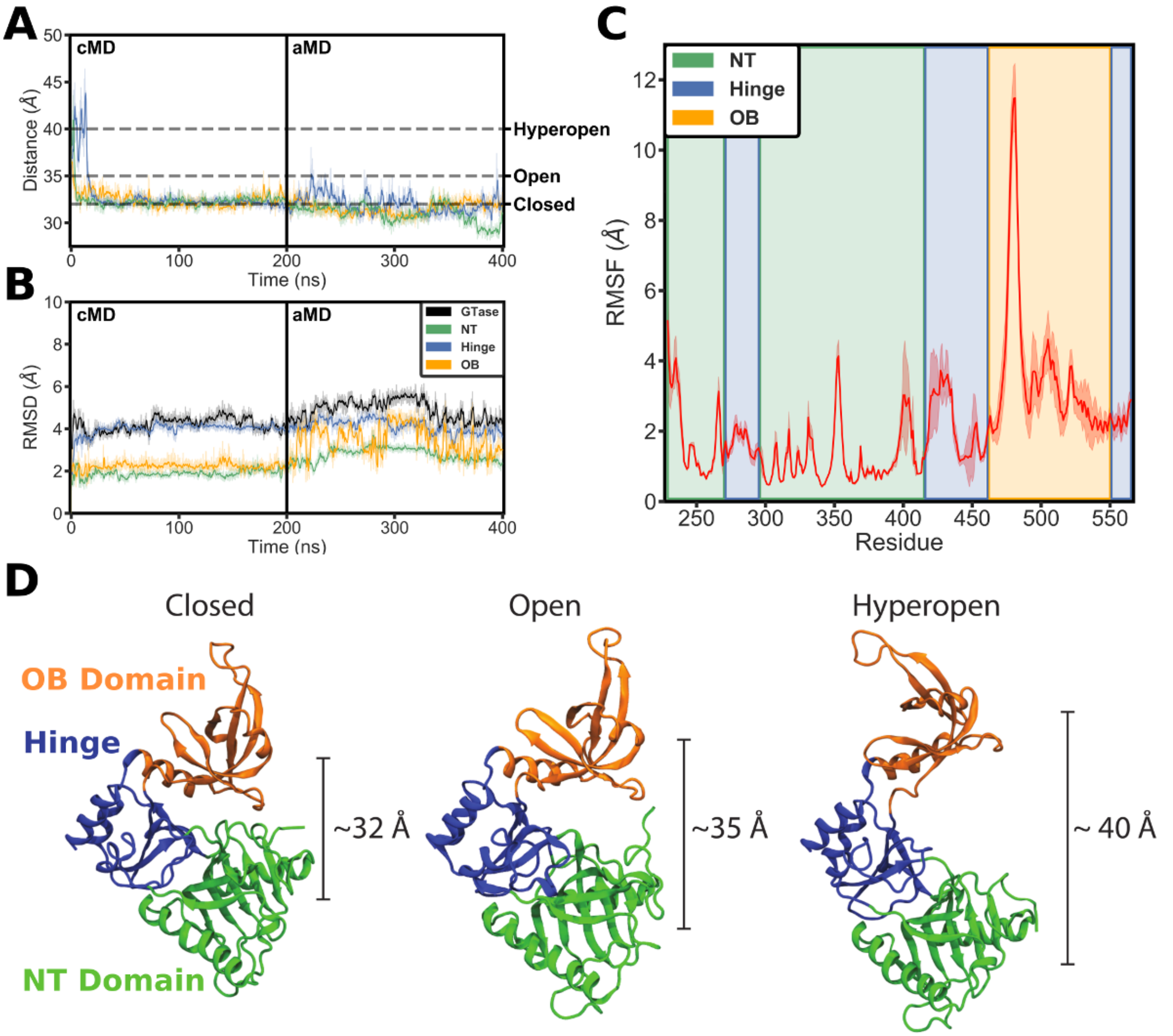


**Figure S1.** Global conformational dynamics of the *apo* human CE GTase. (**A**) The inter-domain distance (centre-of-mass separation) between the OB domain and NT domain over the duration of the cMD and aMD simulations. Three replicates are shown in orange (replicate 1), blue (replicate 2) and green (replicate 3). (**B**) Backbone RMSDs of the whole GTase domain and its sub-domains during the cMD and aMD trajectories. The results are shown for replicate 1; similar results were obtained for two other replicates. The first frame of the cMD was used as a reference. (**C**) Backbone RMSFs of the whole GTase domain obtained from aMD simulations. RMSF values represent the mean of the three aMD replicates. The NT domain in the first frame of the cMD was used as a reference. The shaded area represents one standard deviation. Sub-domains of the GTase structure are highlighted by coloured regions. (**D**) The conformational states that can be adopted by the *apo* CE GTase are designated as ‘Closed’ (separation between the centres of mass of the OB and NT domains ~32 Å), ‘Open’ (~35 Å) and ‘Hyperopen’ (>39 Å), in line with the definitions for the PBCV1 GTase by Swift *et al.* (1).

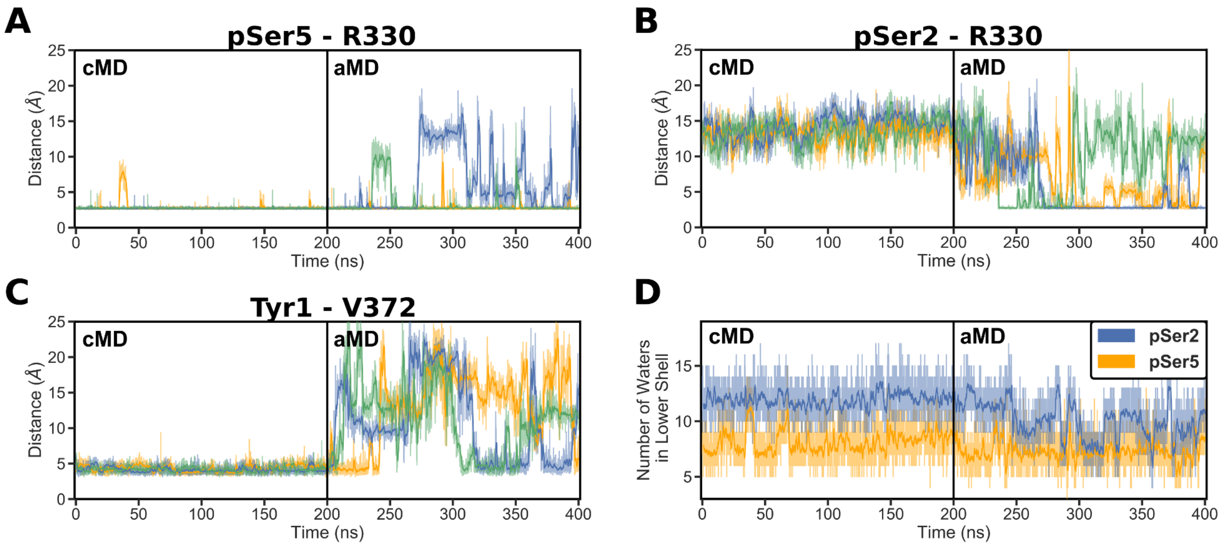


**Figure S2.** Characterisation of the GTase-CTD interaction sites in the 1-heptad CTD simulation (Ser2 and Ser5 phosphorylated; System 3). (**A-B**) Time-evolution of the minimum distances between, CDS1 (taken as the R330 sidechain nitrogens) and pSer5 (A) or pSer2 (B) of the CTD (taken as the phosphate oxygens). (**C**) Time-evolution of the minimum distance between CDS-Y1 (V372 C$\gamma$ atoms) and Tyr1 (sidechain ring) of the CTD. Replicates in (A-C) are displayed in orange (replicate 1), blue (replicate 2) and green (replicate 3). (**D**) Solvent exposure of the pSer2 and pSer5 groups in replicate 1. Number of waters in the lower solvation shell ($<$ 3.4 Å) of the phosphate group is plotted over the cMD and aMD simulation trajectories.

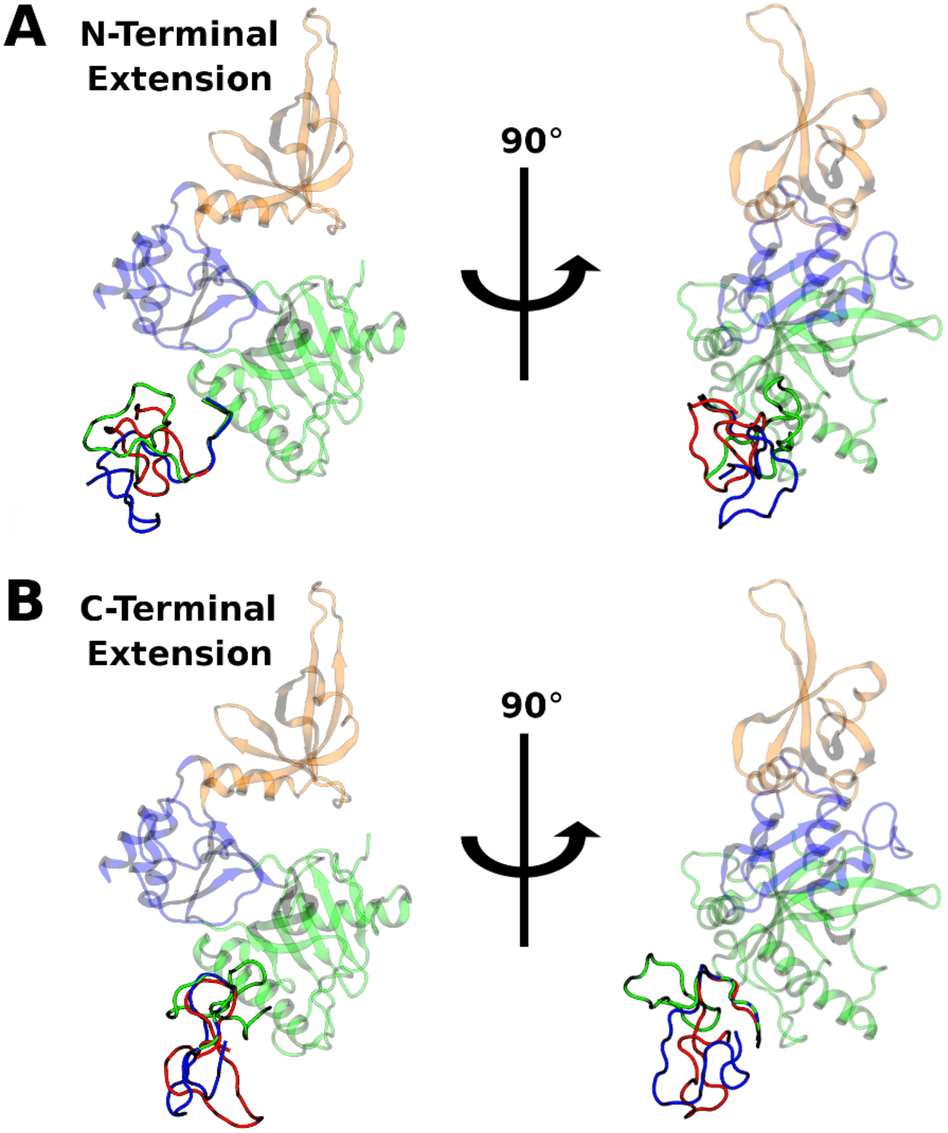


**Figure S3.** Starting CTD conformations in the simulations with a 4-heptad CTD. Each of the three replicates used a different starting structure of the CTD peptide, which were generated using the PEPFOLD3.5 server (2), extending the CTD either in the N-ter direction (**A**) or in the C-ter direction (**B**) (see details in the Methods). The CTD conformations in the three replicates are shown in red (replicate 1), blue (replicate 2) and green (replicate 3).

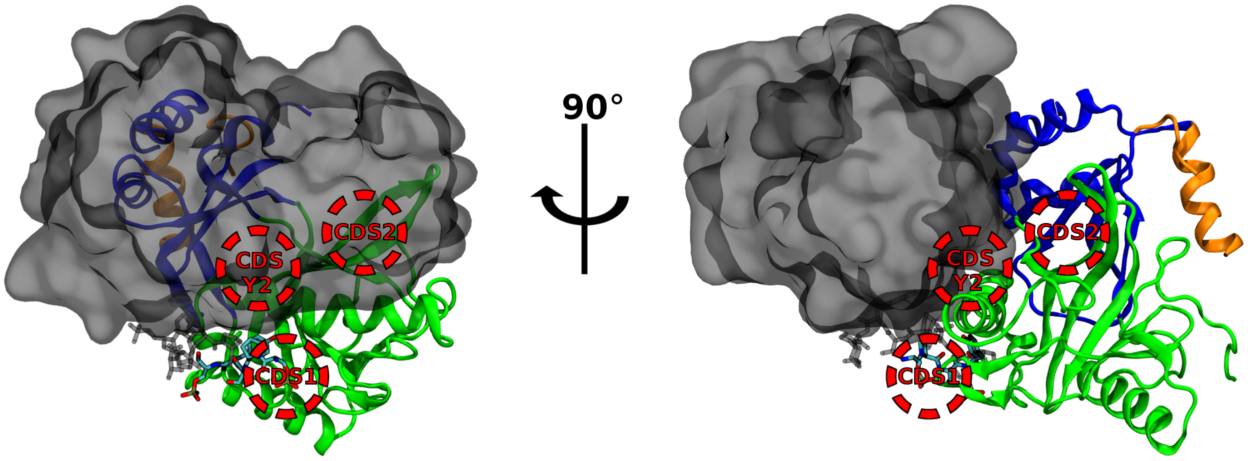


**Figure S4.** Obstruction of CDS2 and CDS-Y2 sites in the previously reported cocrystal structure of the mouse CE GTase-CTD complex (PDB ID: 3RTX) (3). CTD access to the CDS2 and CDS-Y2 sites is blocked by the artificial homodimer interface in the asymmetric unit. The first monomer of the GTase is displayed in the cartoon representation, the second monomer is displayed with a transparent grey surface, and the CTD is shown in stick representation. CDS1, CDS2 and CDS-Y2 locations are indicated in red.

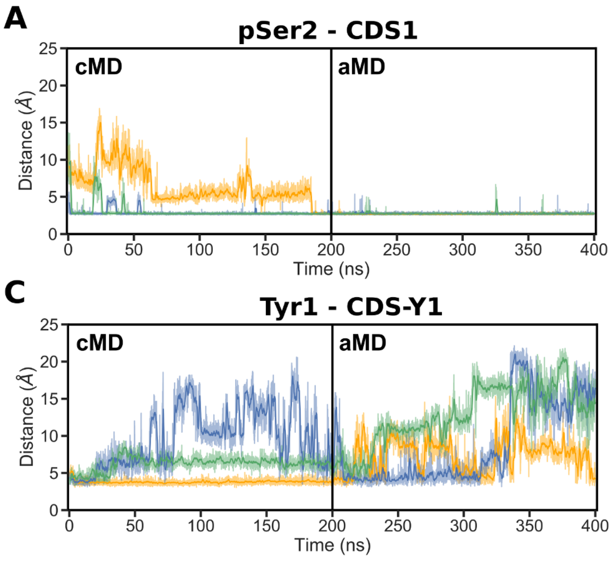


**Figure S5.** Time-evolution minimum distances showing the occupation of each CDS site by the respective pSer2 CTD group over the duration of the cMD and aMD simulations of System 9 (4-heptad, pSer2 CTD extended in the C-ter direction). Distances obtained in three replicates shown as orange (replicate 1), blue (replicate 2) and green (replicate 3). The occupation of each site was described by taking representative sidechain minimum distances as follows: (**A**) CDS1, R330 sidechain nitrogens to the pSer2 phosphate oxygens, (**B**) CDS2, R411 sidechain nitrogens to the pSer2 phosphate oxygens, (**C**) CDS-Y1, V372 C$\gamma$ atoms to the Tyr1 ring, and (**D**) CDS-Y2, L381 sidechain to the Tyr1 ring.

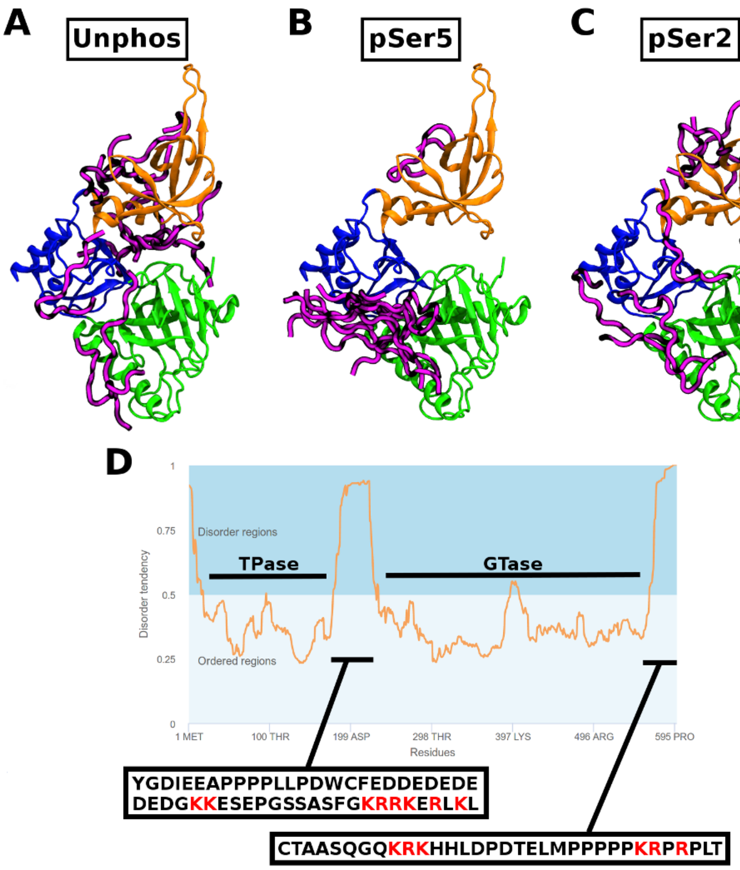


**Figure S6.** Exploring potential CTD interaction sites in regions distal from the NT domain. (**A-C**) Top 10 ranking models predicted by the PIPER-FlexPepDock server showing the binding conformations of the 2-heptad CTD (magenta) in the unphosphorylated (A), Ser5 phosphorylated (B), and Ser2 phosphorylated (C) states. Glutamic acid was used as a phosphomimetic. In the unphosphorylated state, the CTD peptide forms interactions with many regions of the GTase (likely unspecific). The pSer5 phosphorylated peptide docks specifically to the NT domain in the region sampled during our MD simulations, indicating the preferred localisation of pSer5 CTD. The PIPER-FlexPepDock method also identified pSer5 interactions with the novel CDS2 interaction site in 4 of the top 10 models. The pSer2 peptide was seen docked to the CDS2 site in 2 of the top 10 models, but in general can dock into a much wider number of regions on the GTase compared to the pSer5 CTD. (**D**) Disorder prediction for the whole human Capping Enzyme sequence as obtained from the MetadisorderMD2 server (4). Regions predicted to be disordered can be seen at both the N- and C-terminal sides of the GTase domain (these were not resolved in any of the crystal structures). The amino acid sequences of these regions are displayed in boxes, and positively charged residues are highlighted in red.

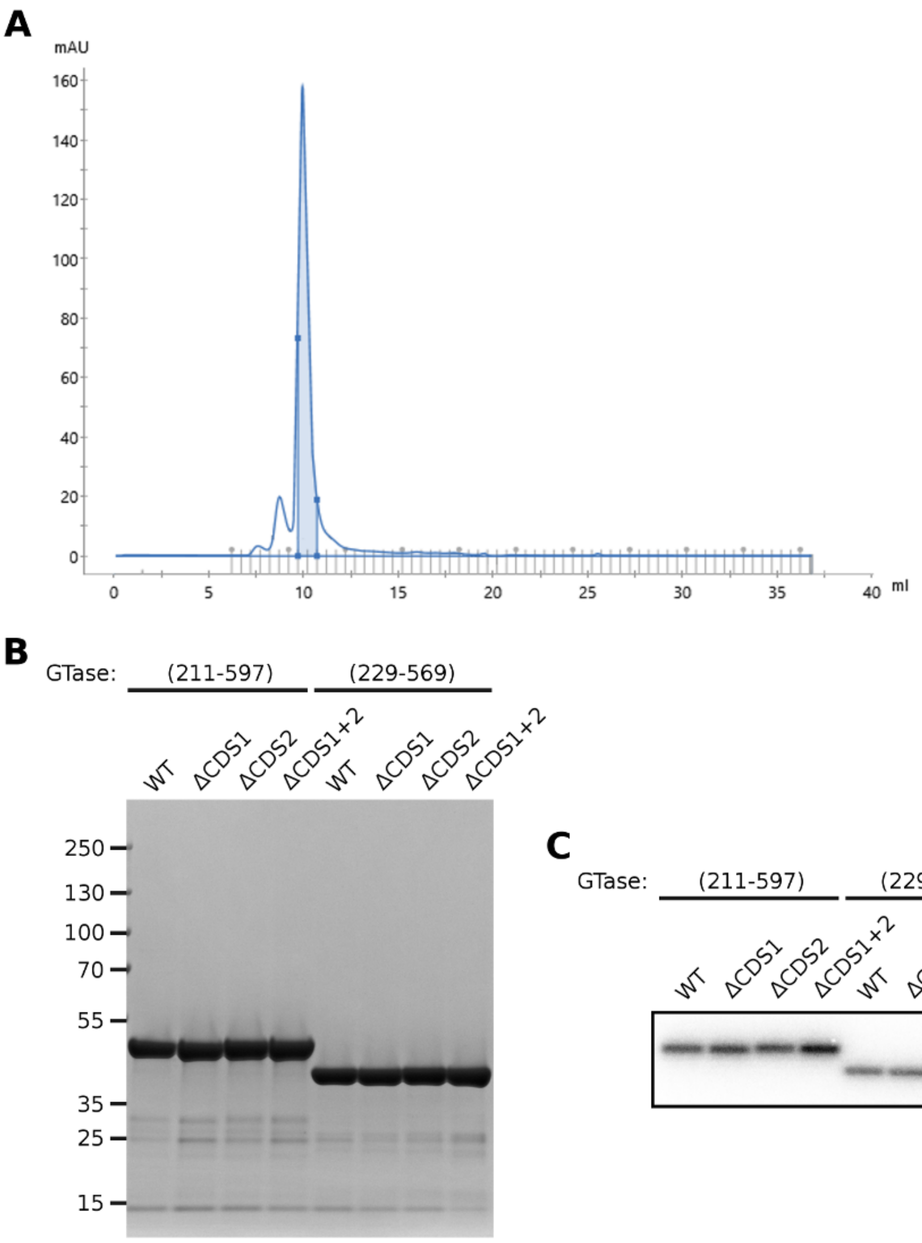


**Figure S7.** Purity and basal activity of the human CE GTase constructs. (**A**) UV chromatograph of the size-exclusion chromatography stage of purification showing a representative elution profile (WT human CE GTase, residues 211-597), with the pooled fractions highlighted in blue. (**B**) Coomassie-stained SDS-PAGE gel of all constructs after the purification process. (**C**) Basal guanylyltransferase activity assay of all constructs.

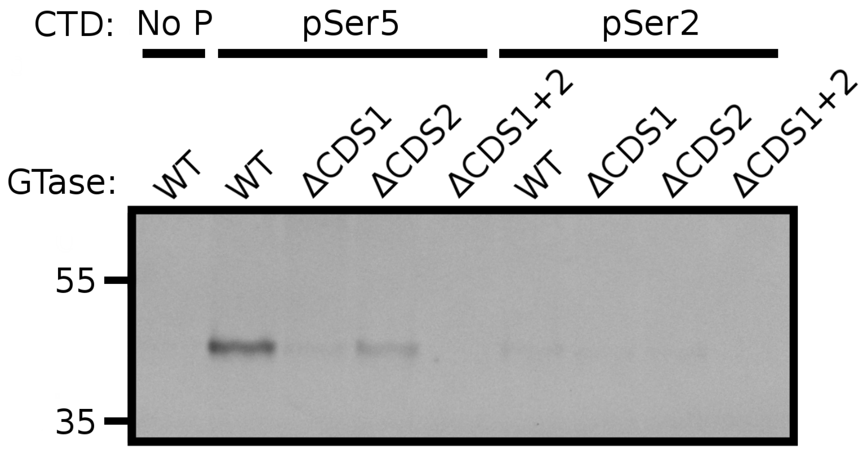


**Figure S8.** Pull-down assay testing the CTD binding affinity in the wild-type and mutant GTase (229-569) for the unphosphorylated, pSer5 and pSer2 CTD peptides. Recombinant human CE GTase was incubated with biotinylated peptides of 4 CTD heptads, in either their unphosphorylated (no P CTD) or phosphorylated state (pSer5 or pSer2 CTD) bound to streptavidin-coupled Dynabeads. The level of GTase binding to the CTD peptides was assessed by SDS-PAGE stained with Coomassie Blue.

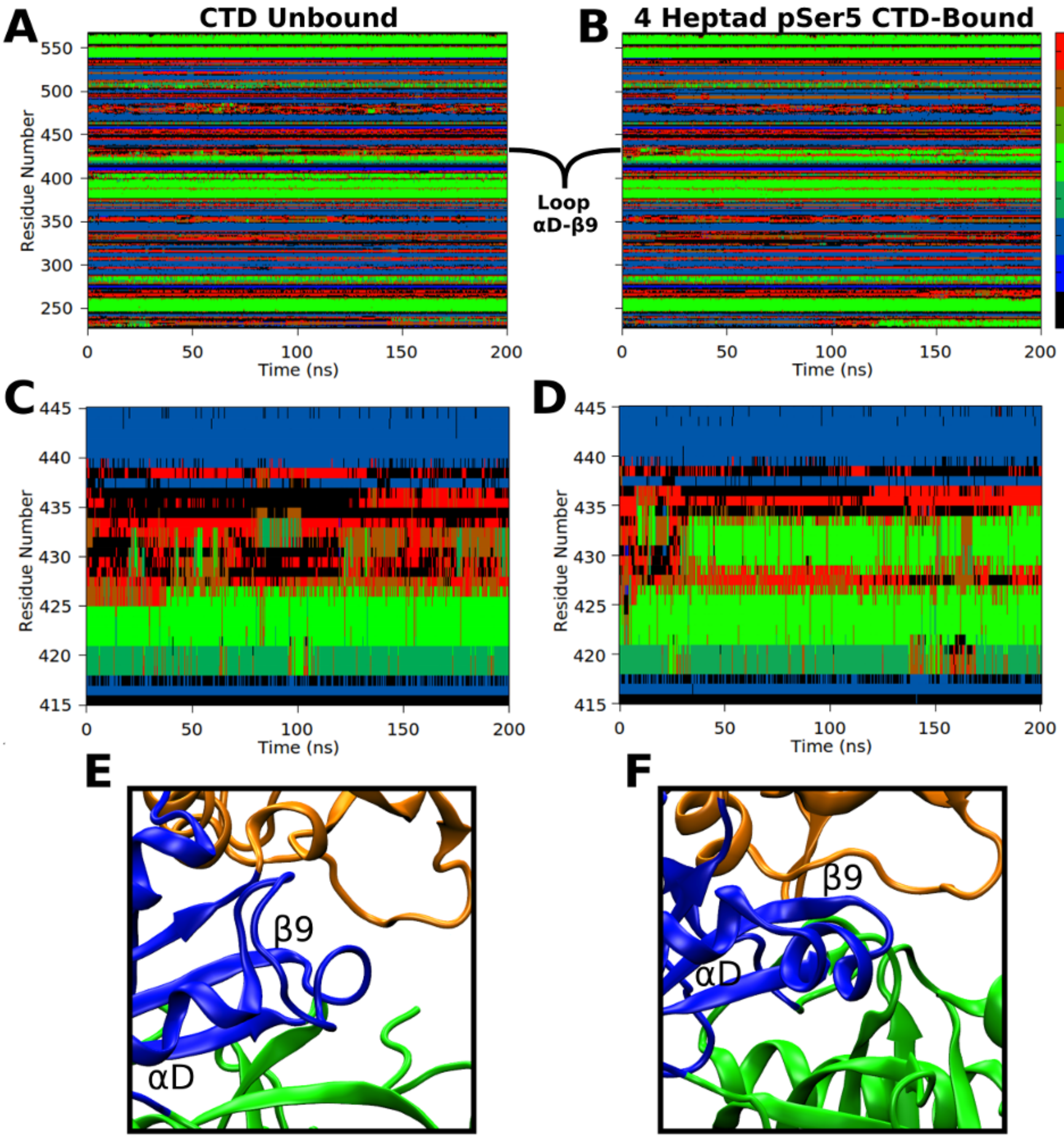


**Figure S9.** Comparing the GTase conformational dynamics between the GTase in the absence of CTD and the 4-heptad pSer5 CTD-bound (N-ter extended) aMD simulations (Systems 1 and 6). (**A-D**) DSSP (Define Secondary Structure of Protein) analysis of the no-CTD system (left, A and C) and 4-heptad pSer5 CTD-bound system (right, B and D), showing the full GTase domain (top, A and B) and loop $\alpha$D-$\beta$9 (middle C and D). (**E-F**) aMD representative snapshots showing a conformation adopted by the loop $\alpha$D-$\beta$9 in the no-CTD system (E) and the pSer5 CTD-bound system (F).

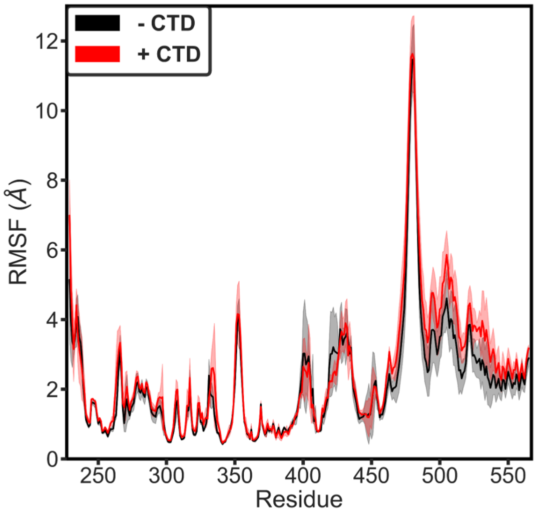


**Figure S10.** Conformational dynamics of the GTase in the presence and absence of the CTD. Backbone RMSFs were calculated for the no-CTD (System 1, **black**) and four heptad pSer5 CTD-bound (N-ter extended; System 6; **red**) GTase systems. RMSF values represent the mean of the three aMD replicates. The NT domain in the first frame of the cMD was used as a reference. The shaded area around the curve represents one standard deviation.

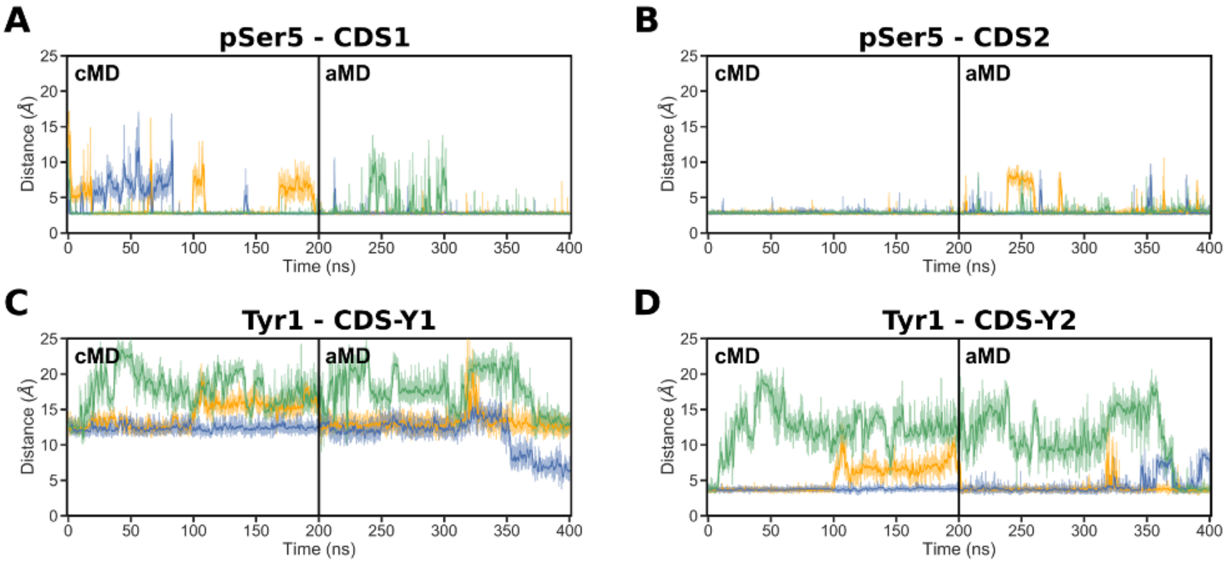


**Figure S11.** The GTase-CTD interaction in simulations started with the pSer5 CTD in the *C. albicans* starting conformation (the 'yeast' orientation, System 10). (**A-D**) Time-evolution of the minimum distances showing the occupation of each CDS by the respective pSer5 CTD group over the duration of the cMD and aMD simulations. The distances are defined as in Figure 3 in the main text. Distances obtained in three replicates are shown in orange (replicate 1), blue (replicate 2) and green (replicate 3). The distance between the CDS1 and pSer5 phosphate in the *C. albicans* GTase-CTD cocrystal structure is only ~11.5 Å. One pSer5 group of the CTD was seen to occupy the 'mammalian' CDS1 site within 20 ns of the simulations and this interaction remained stable. This suggests that the locations of the 'mammalian' and 'yeast' CDS1 sites are close enough on the GTase surface to perform the same underlying role of CTD binding. The CDS2 site remains occupied for the duration of the simulations.

### Movies

**Movie S1** — Overview of the CTD interaction sites observed during the simulations of the GTase with the 4-heptad pSer5 CTD extended in the N-ter direction (System 6, replicate 1).

**Movie S2** — pSer5 binding and the residues involved in the CDS2 interaction site.

**Movie S3** — Residues involved in the CDS-Y2 interaction site.

### PDBs

Conformations of the GTase-CTD systems at the final snapshots of the corresponding simulation. The protein/peptide coordinates are given in the PDB format.

### 1 References


1. Swift, R. V. & McCammon, J. A. Substrate Induced Population Shifts and Stochastic Gating in the PBCV-1 mRNA Capping Enzyme. *Journal of the American Chemical Society* **131**, 5126–5133 (2009).

2. Lamiable, A. *et al.* PEP-FOLD3: faster de novo structure prediction for linear peptides in solution and in complex. *Nucleic Acids Research* **44**, W449–W454 (2016).

3. Ghosh, A., Shuman, S. & Lima, C. D. Structural Insights to How Mammalian Capping Enzyme Reads the CTD Code. *Molecular Cell* **43**, 299–310 (2011).

4. Kozlowski, L. P. & Bujnicki, J. M. MetaDisorder: a meta-server for the prediction of intrinsic disorder in proteins. *BMC Bioinformatics* **13**, 111 (2012).
